# Distinct responses to rare codons in select *Drosophila* tissues

**DOI:** 10.1101/2022.01.06.475284

**Authors:** Scott R. Allen, Rebeccah K. Stewart, Michael Rogers, Ivan Jimenez Ruiz, Erez Cohen, Alain Laederach, Christopher M. Counter, Jessica K. Sawyer, Donald T. Fox

## Abstract

Codon usage bias has long been appreciated to influence protein production. Yet, relatively few studies have analyzed the impacts of codon usage on tissue-specific mRNA and protein expression. Here, we use codon-modified reporters to perform an organism-wide screen in *Drosophila melanogaster* for distinct tissue responses to codon usage bias. These reporters reveal a cliff-like decline of protein expression near the limit of rare codon usage in endogenously expressed *Drosophila* genes. Near the edge of this limit, however, we find the testis and brain are uniquely capable of expressing rare codon-enriched reporters. We define a new metric of tissue-specific codon usage, the tissue-apparent Codon Adaptation Index, to reveal a conserved enrichment for rare codon usage in the endogenously expressed genes of both *Drosophila* and human testis. We further demonstrate a role for rare codons in restricting protein expression of an evolutionarily young gene, *RpL10Aa*, to the *Drosophila* testis. Rare codon-mediated restriction of this testis-specific protein is critical for female fertility. Our work highlights distinct responses to rarely used codons in select tissues, revealing a critical role for codon bias in tissue biology.

## INTRODUCTION

The genetic code is redundant, with 61 codons encoding only 20 amino acids (Crick, 1968; Zuckerkandl & Pauling, 1965). It was initially thought that synonymous substitutions, those leading to changes in nucleotide sequence but resulting in an identical protein sequence, were functionally “silent.” However, it is now clear that this is not the case. Synonymous codons are used at varying frequencies throughout a given genome (Grantham et al., 1980; Ikemura, 1985; Sharp & Li, 1986). This disproportionate usage frequency among synonymous codons is termed codon usage bias (hereafter: codon bias).

Codon bias is governed by several biochemical mechanisms and has diverse biological consequences. In general, mRNAs enriched in codons commonly used in a given species are more stable and are more robustly translated (Presnyak et al., 2015; Sørensen & Pedersen, 1991; Yan et al., 2016; Yu et al., 2015; Zhao et al., 2017). Conversely, a high frequency of rare codons in an mRNA can cause ribosomal stalling and trigger RNA degradation (Buschauer et al., 2020; Radhakrishnan et al., 2016). Codon bias impacts protein expression that underlies circadian regulation of growth in *Neurospora* (Xu et al., 2013; M. Zhou et al., 2013), protein secretion in yeast (Pechmann et al., 2014), and virus/host interactions (including in COVID-19, Alonso & Diambra, 2020; Shin et al., 2015).

Emerging studies suggest codon bias plays an important role in fundamental tissue-level processes. For example, codon bias underlies differences between maternal and zygotic mRNAs in developing zebrafish, *Xenopus*, mouse, and *Drosophila* (Bazzini et al., 2016). Further, while rare codons generally destabilize mRNAs in *Drosophila* whole embryos, this effect is attenuated within the embryonic central nervous system, where codon bias has little impact on mRNA stability (Burow et al., 2018). Studies analyzing gene expression datasets cross-referenced with codon usage suggest that the impact of codon bias on protein expression may differ between tissues. For example, codon usage frequencies differ between tissue-specific gene sets in numerous plant species (Camiolo et al., 2012; Liu, 2012), *Drosophila* (Payne & Alvarez-Ponce, 2019), and potentially humans (Plotkin et al., 2004; Sémon et al., 2006), hinting that codon usage could play a fundamental role in tissue and cellular identity.

Here, we leverage the genetic and cell biological strengths of *Drosophila melanogaster* to reveal tissue-specific impacts of codon bias. Using a library of codon-altered reporters, we conduct an organism-wide screen during *Drosophila* development. We find that reporter protein expression declines drastically over a narrow range of rare codon usage in a gene. We further show that specific tissues, namely the testis and brain, are distinct in the ability to robustly express proteins encoded by rare codons. Focusing further on the testis, the tissue with the strongest protein expression from rare codon-enriched genes, we find the male germ cells and somatic hub cells are capable of robust rare codon-derived protein production, while somatic cyst cells are not. By developing a new metric to examine tissue-specific codon usage, tissue-apparent CAI (taCAI), we find that both *Drosophila* and human testes express genes that are enriched in rare codons relative to other tissues. Examining the physiological significance of this conserved enrichment, we highlight a role for rare codons in restricting protein expression of RpL10Aa, a tissue-specific ribosomal subunit and evolutionarily young gene, to the testis. This rare codon-imposed restriction is critical for fertility. Just as chromatin states regulate tissue-specific gene expression at the level of transcription, here we find a biologically significant role for codon bias, an important determinant of mRNA translation, in regulating tissue-specific protein production.

## RESULTS

### A cliff-like protein expression threshold determined by rare codons in *Drosophila*

We previously showed that severely altering codon content impacts protein expression from *RAS* family genes or from a *GFP* reporter in *Drosophila melanogaster* and humans (Ali et al., 2017; Lampson et al., 2013; Peterson et al., 2020; Sawyer et al., 2020). This prompted us to systematically determine the impact of codon usage on protein expression. To do so, we again used phiC31 integrase for stable site-specific insertion of a single copy of transgenic codon-modified *GFP* reporters (**Table S1**) into the *attP40* locus on chromosome 2. This locus is well-established to express transgenes at robust and reproducible levels (Markstein et al., 2008). As we have done previously (Sawyer et al., 2020), we placed all our transgenic reporters in the same vector backbone, which uses a *ubiquitin* promoter sequence for robust gene expression (Methods). In this way, we control for the effects of chromatin environment on gene expression and ensure that any differences in protein production are due to differences in coding sequence.

We took two approaches to introduce rare codons into our *GFP* reporters. In the first approach, we used a random number generator to select positions in the *GFP* coding sequence to engineer rare codons. This approach keeps the majority of rare codons dispersed throughout the *GFP* sequence (**Fig 1A**). We generated 10 reporters using this dispersed codon strategy, and such reporters are designated with a “D”. Given the potential importance of contiguous stretches of rare codons (Chu et al., 2014; Hayes et al., 2002; Kramer & Farabaugh, 2007; Spanjaard & van Duin, 1988), we also employed a second approach in which all rare codons were clustered at either the 5’ or 3’ end of the coding sequence. We generated 6 reporters using this clustered codon strategy, and all reporters generated this way are designated “C5’” or “C3’”. In defining rare codons, we referred to the Kazusa codon database (Nakamura et al., 2000). We identified the most used synonymous codon in the *Drosophila melanogaster* genome as the “common” codon, the least used codon as the “rare” codon, and any additional synonymous codons between the rare and common codon as “middle” codons. We designed reporters to contain a specific percentage of rare codons, leaving the remaining portion of the coding sequence split between common and middle codons (**Fig 1A**). These reporters were all designed using an identical eGFP amino acid sequence to rule out any effects of amino acid bias on protein production levels (Weber et al., 2020). All reporters are designated with the fluorophore (e.g. GFP) followed by the percentage of rare codons, followed by a “D” or “C” designation (e.g. *GFP50D*, *GFP70C3’*).

**Figure 1.**
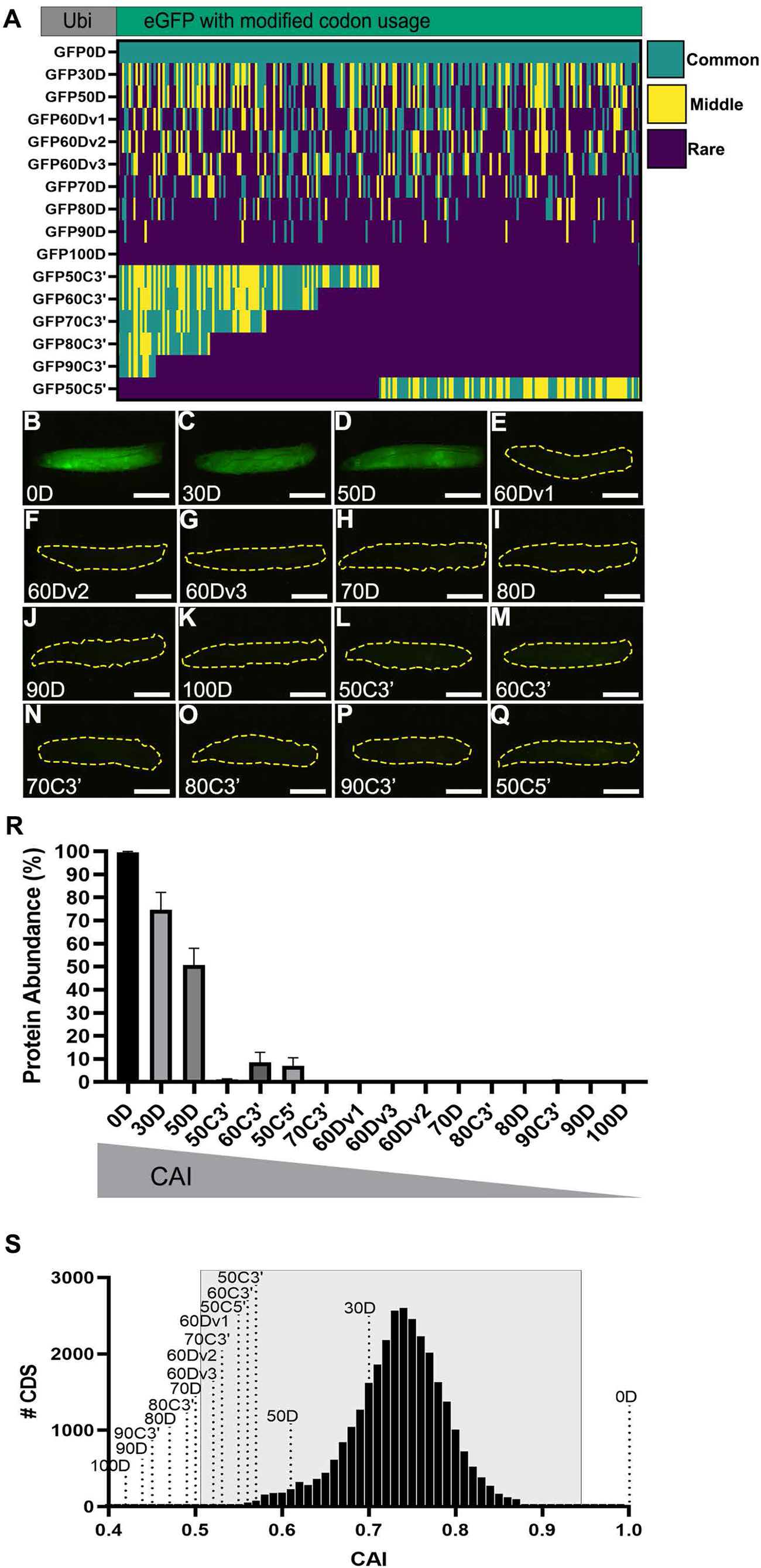
Strictly defined limits on rare codon usage regulate protein production. (**A**) Top-schematic of each indicated reporter. Bottom-heat map indicating codon usage across each reporter coding sequence. Color key indicates codon usage along the length of the coding sequence. (**B-Q**) Representative fluorescent images taken under identical camera exposure settings of live male WL3 larvae containing stable genomic insertions of the indicated *GFP* reporter. Scalebars are 1mm. Yellow dashed outlines highlight the larvae in images where no GFP signal was apparent. (**R**) Western blot protein quantifications of indicated reporters normalized to total protein stain then plotted as percentages relative to GFP0D for whole third instar larvae (2 replicates, N=5 each, plotting mean +/-SEM). Reporters listed in descending order by CAI. See **Figure1-Figure Supplement 1** for blot images. (**S**) Histogram of CAI values for each transcript in the *Drosophila melanogaster* genome. Reporter CAI values are indicated with dotted lines. Full range of endogenous genes highlighted in light gray box.

As a convenient method to screen how codon substitutions in GFP impact protein expression across the entire animal, we imaged wandering third instar larvae (WL3) for each transgenic reporter on a fluorescent dissection microscope. These animals contain fully formed organ systems and are translucent to facilitate imaging. We used the *GFP0D* reporter with no rare codons as a baseline for comparison. Upon examining animals for each reporter, we noticed an apparent “all or none” fluorescence pattern (**Fig 1B-Q**). To quantitate our observations seen by fluorescent microscope, we measured protein production by western blot (**Fig 1R, Figure 1-Figure Supplement 1**). For reporters with dispersed rare codons, 2/2 reporters with up to 50 percent rare codons display ubiquitous fluorescence throughout the animal, similar to *GFP0D*. These two reporters (*GFP30D* and *GFP50D*) produce animal-wide GFP protein at levels greater than or equal to 50 percent of GFP0D (**Fig 1B-D,R, Figure 1-Figure Supplement 1A**). In contrast, 7/7 reporters with greater than 50 percent dispersed rare codons display no visible fluorescence in the animal and produce protein at a level less than 0.3 percent of GFP0D (**Fig 1B,E-K,R, Figure 1-Figure Supplement 1B,C**). Similarly, 6/6 reporters with 50 percent or more clustered rare codons display no fluorescence (**Fig 1L-Q**). Two of these six clustered codon reporters, *GFP50C5’* and *GFP60C3’*, produce barely detectable protein levels by western blot, between 7-9 percent that of GFP0D, whereas the other four reporters yield less protein than 1 percent of GFP0D (**Fig 1R, Figure 1-Figure Supplement 1C,D**). While it is not surprising that there would be a rare codon threshold beyond which proteins are no longer produced, what is surprising is the magnitude of the differences in protein production (from robust to barely/not detectable) over a narrow range of rare codon usage. A 50 percent increase in rare codon usage between *GFP0D* and *GFP50D* causes a 50 percent reduction in protein levels. Yet an additional 10 percent increase in rare codon usage between *GFP50D* and *GFP60Dv1* decreases detectable protein by 99.5 percent (**Fig 1R, Figure 1-Figure Supplement 1A**).

To determine if the steep decline in protein levels between 50 and 60 percent dispersed rare codons is robust and reproducible, we independently designed two more 60 percent dispersed rare codon reporters. These reporters, *GFP60Dv2* and *GFP60Dv3*, contain rare codons that differ in sequence/position from *GFP60Dv1*. All three GFP60D reporters show the same fluorescence pattern (no detectable GFP), and when protein levels are compared to GFP50D by western blot we again observe a decrease in protein levels of at least 99.5 percent, with a 99.8 percent decrease on average across all three GFP60D reporters (**Fig 1E-G,R, Figure 1-Figure Supplement 1B**). Overall, our results indicate there is a narrowly defined window of rare codon content in which protein production drops from robust to barely detectable/none.

We next put our observed cliff-like decline in protein levels from reporters in context with endogenous *Drosophila melanogaster* gene sequences. To look at the codon usage of endogenous genes, we turned to the well-established Codon Adaptation Index (CAI) as a metric for measuring codon usage optimality (Sharp & Li, 1986). CAI calculates the degree of optimal codon usage in a transcript based on the usage frequency of that codon in a reference set of genes. Higher CAI scores indicate more common codon usage, where a score of 1 indicates a transcript only uses the most common codons for each amino acid (Nakamura et al., 2000). We plotted the CAI values of each individual transcript in the *Drosophila melanogaster* genome (**Fig 1S**, Methods). We similarly computed the CAI values for each of our codon-modified *GFP* reporters (**Fig 1 R,S**).

Comparing CAI values between reporters and endogenous genes reveals a striking cliff-like protein expression limit imparted by rare codons. This limit is reflected in the all-or-none expression pattern of our reporters. The lowest CAI of any reporter (*GFP50D*, **Fig 1R, Figure 1-Figure Supplement 1**) with detectable fluorescence resides near the limit of CAI for endogenously expressed genes (only above the CAI of 2% of all expressed genes-**Fig 1S, see left boundary of gray rectangle**). Below the CAI value of *GFP50D*, none (0/13) of our reporters are expressed at the level of detectable fluorescence, and only two of these reporters produce protein detectable by western blot at low levels (**Fig 1R, Fig 1-Fig Supplement 1**). Our findings suggest that in *Drosophila,* a limit on robust rare codon-derived protein expression is in the range of 50 to 60 percent rare codon usage or between CAI scores of 0.61 and 0.57.

### The testis and brain robustly express rare codon-enriched reporters

While overall we observe a cliff-like threshold in reporter GFP levels (**Fig 1R, Figure 1-Figure Supplement 1**), none of the clustered rare codon reporters reside within our determined CAI-dependent limit to be detected by fluorescence. Given this, we generated another clustered codon reporter (*GFP54C3’*). In designing *GFP54C3’*, we clustered all of the rare codons in the 3’ end of the coding sequence and used only the most common or most rare codons for each amino acid. Compared to *GFP50D,* this design increased rare codon content while maintaining a higher overall CAI score (0.64, **Fig 2A**). *GFP54C3’* falls near the limit of CAI for endogenously expressed *Drosophila* genes, ranking higher than just 5 percent of endogenous transcripts. Unlike the all-or-none expression pattern of the reporters shown in **Fig 1**, GFP54C3’ shows no signal in most of the animal but has a robust GFP fluorescence in two distinct areas of the larva (**Fig 2C**). This suggests that distinct tissues respond differently to rare codons.

**Figure 2.**
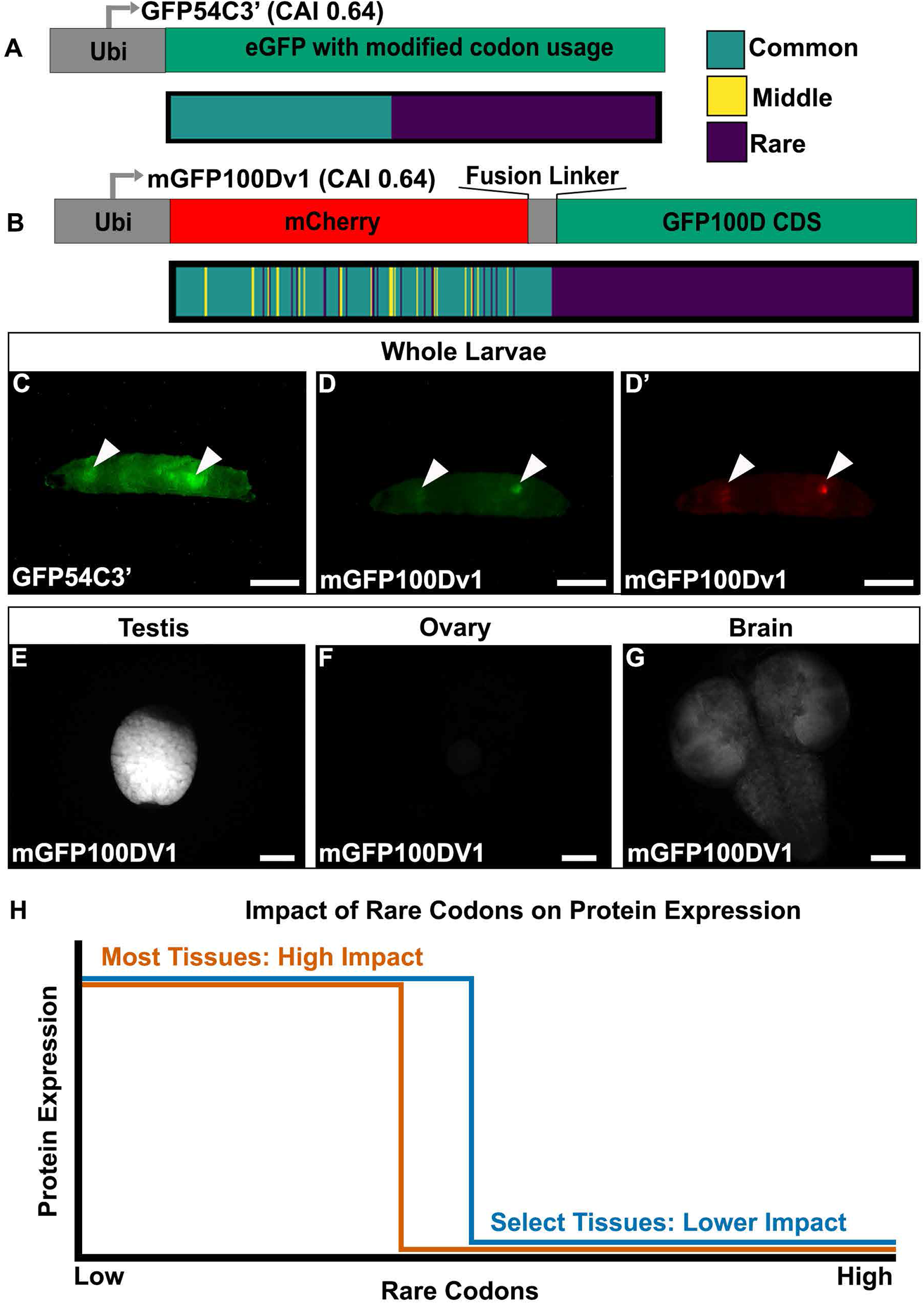
Tissues exhibit distinct responses to rare codons. (**A**) Top-reporter design and Bottom-codon usage (see key) along the CDS of *GFP54C3’*. CAI for *GFP54C3’* indicated in parentheses. (**B**) Top-reporter design and Bottom-codon usage along the CDS of *mGFP100Dv1* (see key in panel A). CAI for *mGFP100Dv1* indicated in parentheses. (**C**) Representative fluorescent images of live male WL3 larva with stable genomic insertion of *GFP54C3’*. (**D**) Representative fluorescent image of live male WL3 larva with stable genomic insertion of *mGFP100Dv1* taken at GFP excitation wavelength. (**D’**) Representative fluorescent image of live male WL3 larva with stable genomic insertion of *mGFP100Dv1* taken at mCherry excitation wavelength. Scalebars for **C-D’** are 1mm. (**E-G**) Representative fluorescent images of dissected *mGFP100Dv1* larval tissues. Images are taken under identical conditions with fluorescence intensity normalized to testis. Scalebars for **E-F** are 100um. (**H**) Conceptual depiction of tissue specific differences in rare codon tolerance, as revealed by our reporters. As rare codons increase (moving to the right on the X-axis), protein expression (Y-axis) undergoes a cliff-like decline. However, the point at which select tissues hit the edge of the cliff is distinct.

Given the unique tissue-specific pattern with this reporter, we generated additional reporters to further understand the sequence parameters driving this expression. To assess whether the tissue-specific expression of GFP54C3’ is due to an unknown sequence motif, we generated an entirely different reporter with similar sequence parameters. To do so, we generated a hybrid *mCherry/GFP* reporter. Both *mCherry* and *GFP* are similar in size, (708 and 717 nucleotides, respectively). Since the rare codons in *GFP54C3’* are clustered towards the 3’ end, we designed *mGFP100Dv1*, where we redesigned *GFP100D* by creating a fusion protein linked to a 5’ *mCherry* sequence (Methods). This *mCherry* is naturally enriched in common codons, and the fusion to *GFP100D* creates a reporter with a CAI similar to *GFP54C3’* (**Fig 2B**). mGFP100Dv1 rescues the lack of expression seen with GFP100D in a tissue-specific manner (**Fig 2D** **vs.** **Fig 1K**). Using further altered versions of this hybrid mCherry/GFP reporter (**Fig 2-Fig Supplement 1A**, Methods), we ruled out that the tissue-specific rescue of expression is due to an unknown sequence motif in the mCherry or fusion linker (**Fig 2-Fig Supplement 1B-G**). We also examined the sequences of all reporters for co-occurring codon pairs known to impact translation fidelity, such as AGG and CGA (Letzring et al., 2010; Spanjaard & van Duin, 1988), and find no predictive pattern in their occurrence between our tissue-specific and non-tissue-specific reporters (**Table S2**).

Having established that tissue-specificity can be achieved using multiple different coding sequences, we next determined which tissues express rare codon-enriched GFP54C3’ and mGFP100Dv1. In larvae, we dissected out the two bright tissues from mGFP100DV1. In doing so, we noticed that male larvae always have two bright tissues, whereas female larvae always have one. Our dissections revealed this difference to be because mGFP100DV1 expresses brightly in the larval testis (**Fig 2E**) but is not detectable in larval ovaries (**Fig 2F**). The other mGFP100DV1 bright tissue, in common to both female and male mGFP100DV1 larvae, is the brain (**Fig 2G**). Together, our results reveal that, within a narrow range of 50-60 percent rare codons, a select few tissues are capable of robustly producing protein from rare codon-enriched transcripts (**Fig 2H**).

We next turned to adult animals, to assess if the distinct testis and brain expression carries forward to this stage. We used a combination of fluorescence microscopy of isolated tissues (**Fig 3A-L’**) and quantitative western blotting (**Fig 3M**, Methods). This analysis again revealed that the testis and brain express mGFP100Dv1 with the testis having the strongest relative protein expression (**Fig 3E,F,M**). Further, we find that other tissues, such as the ovary and male accessory glands, do not express this reporter at a detectable level (**Fig 3G,H,M**). Conversely, the common codon-enriched GFP0D expresses robustly in all adult tissues examined (**Fig 3A-D,M**) suggesting that lack of mGFP100Dv1 expression is not due to differences in promoter strength between tissues.

**Figure 3.**
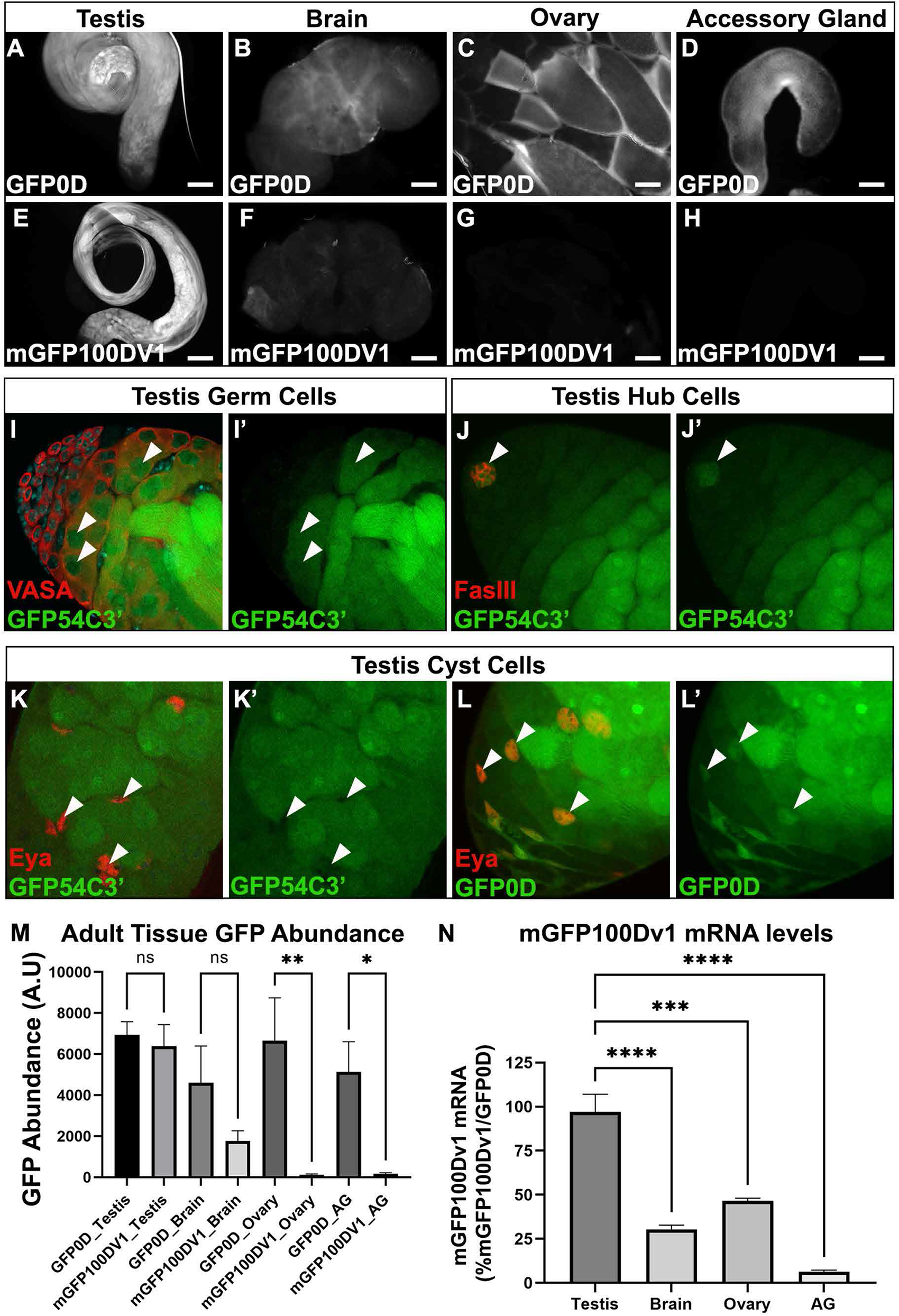
The adult testis and brain robustly express rare codon-enriched reporters. (**A-D**) Representative fluorescent images of dissected *GFP0D* adult tissues. Images are taken under identical conditions with fluorescence intensity normalized to testis. (**E-H**) Representative fluorescent images of dissected *mGFP100Dv1* adult tissues. Images are taken under identical conditions with fluorescence intensity normalized to testis. (**I-L’**) Confocal images of adult *GFP54C3’* testis immunostained with antibodies recognizing the indicated cell types. Arrowheads indicate cells of interest, **I**,**I’**= germ cells, **J,J’**= hub, **K,K’**= somatic cyst cells. Right image in each pair shows the GFP only channel from the left image in the pair. (**L’L’**) Confocal image of adult GFP0D testis immunostained with Eya. Arrowheads= somatic cyst cells. (**L’**) GFP only channel of image in **L** (**M**) Quantitation of GFP protein abundance in dissected adult tissues measured by western blot for *GFP0D* and *mGFP100Dv1* animals normalized to total protein stain (3 replicates, N=10-12 animals each, plotting mean +/-SEM, Bonferroni-Šídák, *p<0.05, **p<0.01). See **Figure 3-Figure Supplement 1** for representative blot image. (**N**) Steady state mRNA levels for heterozygous *mGFP100Dv1/GFP0D* expressing animals measured by qRT-PCR. *mGFP100Dv1* mRNA levels are plotted as a percentage relative to GFP0D within each tissue. (2-3 replicates, N=10 animals each, plotting mean +/-SEM, Dunnett’s multiple comparison to testis, ***p=0.0001,****p<0.0001). Scalebars are 100um.

To quantify the impact of rare codons on protein production in each tissue, we compared protein levels of mGFP100Dv1 to GFP0D from three replicate western blots (**Fig 3M, Fig 3-Fig Supplement 1**). There is no statistically significant difference in protein levels between GFP0D and mGFP100Dv1 for both testis and brain in adult males, consistent with these tissues having a distinctly higher tolerance for rare codons (**Fig 3M**). In contrast, both the ovary and accessory gland produce mGFP100Dv1 protein at levels less than or equal to 2 percent that of GFP0D protein (**Fig 3M**), consistent with a lower rare codon tolerance in these (and most other) *Drosophila* tissues (**Fig 2H**). While we do not observe differences in the ability of the brain to express rare codon-enriched reporters between sexes (**Fig 3-Fig Supplement 2**), the testis but not the ovary robustly expresses protein derived from rare codon-enriched reporters (**Fig 3M**). One possible explanation for the tissue-specific differences in protein expression could be differences in the strength of the *ubiquitin* promoter in each tissue. However, our fluorescence-based approach suggested this is not the case (**Fig 3A-D,M**). We further assessed this by examining the levels of GFP0D (all common codons) by western blot in each tissue. Interestingly, the ovary expresses GFP0D at similar levels to the testis, yet rare codon-enriched mGFP100Dv1 is expressed 100-fold higher in the testis than in the ovary (**Fig 3M**). This confirms that promoter strength is not a confounding variable. Collectively, these results highlight striking differences between *Drosophila* tissues, most notably between the male and female germline, with regards to rare codon-derived protein expression.

Our findings on rare codon-derived protein expression at a whole tissue level, which are most pronounced in the testis, prompted us to further examine which specific cell types in the testis can robustly express rare codon-enriched reporters. Multiple cell types comprise the *Drosophila* testis, including both somatic and germline cell lineages (Fairchild et al., 2017; Herrera & Bach, 2019; Lim et al., 2015; Yamashita et al., 2005). To determine which specific cell types express protein from rare codon-enriched reporters, we used cell type-specific antibodies and assessed co-localization with rare codon-enriched GFP54C3’. Male germ cells (Vasa positive) and the somatic hub cells (FasIII positive), which comprise the male germline stem cell niche, robustly express GFP54C3’ (**Fig 3I-J’**), whereas somatic cyst cells that encapsulate the germline cells (Eya positive) do not (**Fig 3K,K’**). Importantly, somatic cyst cells robustly express common codon-enriched GFP0D (**Fig 3L,L’**), indicating that the *ubiquitin* promoter is active in this cell type. Thus, both the male germ cells and the specialized somatic cells of the male germline stem cell niche express protein derived from a rare codon-enriched reporter, while the larger population of somatic cyst cells lack this capability. Together, we find a striking difference between male and female gonads in the expression of protein from rare codon-enriched genes.

### Distinct regulation of rare codon-enriched mRNAs in the testis of flies and humans

Our observed tissue-specific protein expression of rare codon-enriched reporters in the testis and brain could be primarily driven at the protein level, but could also be regulated at the level of mRNA (reviewed in Radhakrishnan & Green, 2016). Rare codons can decrease levels of transcription (Zhao et al., 2021; Z. Zhou et al., 2016) or can negatively impact mRNA stability through a growing number of characterized mechanisms and in numerous model systems (Bazzini et al., 2016; Burow et al., 2018; Buschauer et al., 2020; Presnyak et al., 2015; Radhakrishnan et al., 2016; Wu et al., 2019). Alternatively, translational repression mechanisms can act independently of mRNA stability, further complicating the relationship between codon optimality and mRNA levels (Freimer et al., 2018).

We next assessed whether reporter protein abundance reflects mRNA levels in different tissues. The highest mRNA level for mGFP100Dv1 is found in the testis (**Fig 3N**). Further, mRNA levels of mGFP100Dv1 and GFP0D are nearly identical in the testis. This result matches our finding at the protein level in the testis (**Fig 3N**). mGFP100Dv1 mRNA is also detected in several other tissues, but at a reduced level relative to GFP0D. Specifically, we detect mGFP100Dv1 mRNA in the ovary (50% of GFP0D levels), brain (30% of GFP0D), and accessory gland (10% of GFP0D, **Fig 3N**). These results show that mRNA levels from a rare codon-enriched reporter differ in various tissues. Among the non-testis tissues examined, only the brain appears able to convert this mRNA into abundant protein. Our findings further contribute to the complex relationship between codon bias, mRNA levels, and protein levels. Further, these mRNA results support our finding that, relative to all other tissues examined, protein expression in the testis is less impacted by codon bias.

To put our findings with synthetic reporter mRNA levels and codon content in context with mRNA levels of endogenous genes, we next examined publicly available tissue-specific *Drosophila* RNAseq datasets (Methods). We developed a modification to the CAI to create a new codon usage metric that reflects codon usage among highly expressed mRNAs that are enriched in a specific tissue. We term this metric tissue-apparent CAI (taCAI). In developing taCAI, we first defined tissue-specific gene sets for each tissue using cutoffs based on relative expression levels (Methods). From these tissue-specific gene sets, we then took the 300 most enriched genes for each of 12 adult tissues and obtained the usage frequency for each codon within that set of over-represented genes. The codon usage frequencies for each coding sequence in the genome were then compared against usage frequencies within the tissue-specific gene set to obtain a spread of taCAI values visualized as violin plots. High taCAI values indicate that mRNAs expressed in a tissue are enriched for rare codons, while low taCAI values indicate that mRNAs expressed in a tissue are enriched for common codons. We calculated taCAI for individual adult tissues in *Drosophila melanogaster* for which RNAseq data exists in FlyAtlas2 (Methods).

When visualizing the taCAI distributions for each tissue as a violin plot (**Fig 4A**), we notice a clear tissue-specific trend. The testis and accessory gland, both parts of the male reproductive system, are highly enriched for mRNAs with abundant rare codon usage relative to all other tissues and to the transcriptome as a whole. We note that the accessory gland does not accumulate rare codon-enriched reporters, and the brain does not have a particularly high taCAI (see **Discussion**). In the testis, our computational findings with taCAI match our experimental results using rare codon-enriched *GFP* reporters. Taken together, our results show that among the tissues examined, the *Drosophila* testis accumulates by far the highest mRNA and protein levels for a rare codon-enriched reporter, and that this finding is consistent with high levels of mRNA for endogenous testis-expressed, rare codon-enriched genes.

**Figure 4.**
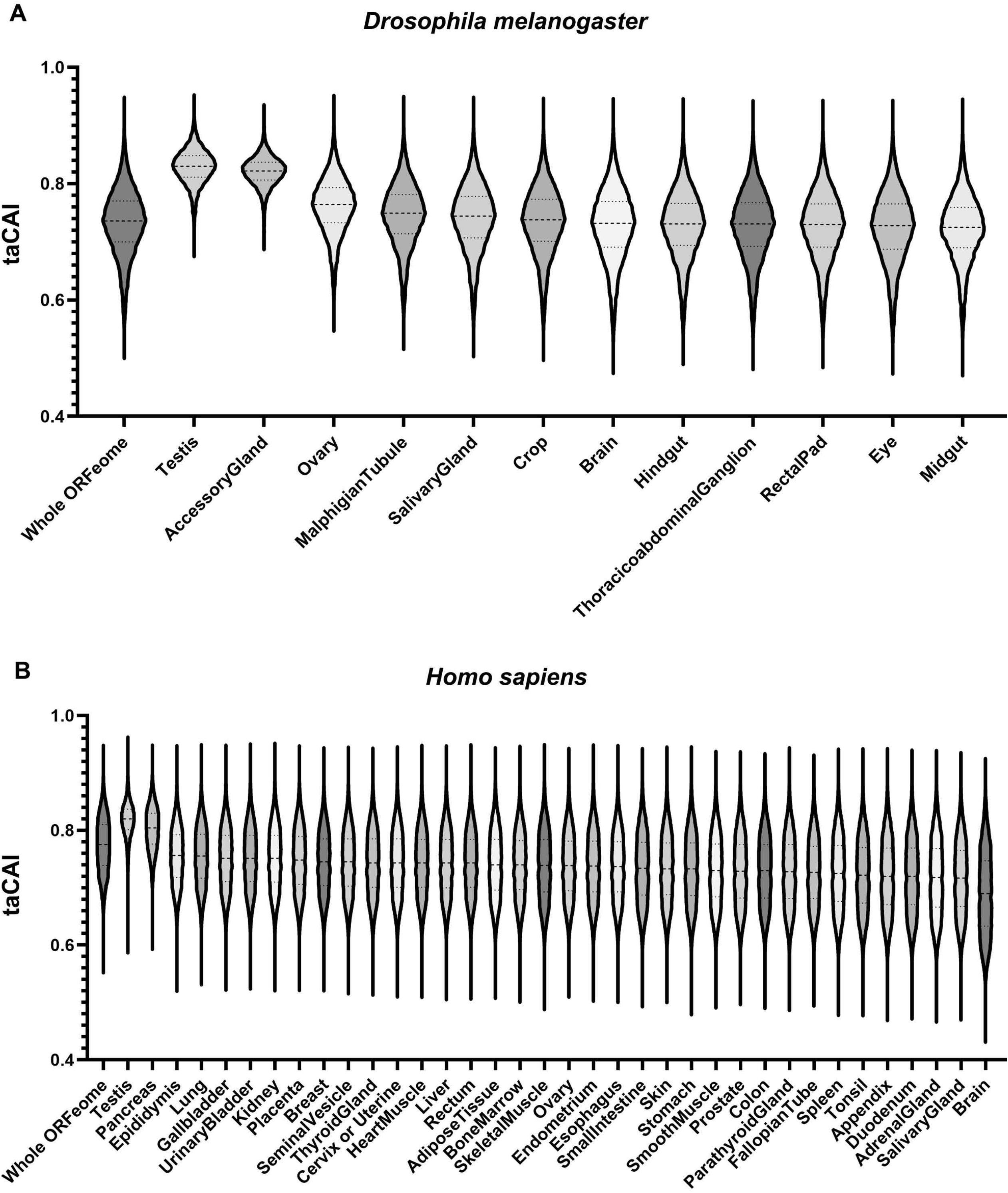
Endogenous testis genes are enriched in rare codons. (**A**) taCAI for *Drosophila melanogaster* tissues from FlyAtlas2. All tissues analyzed are from adult male animals except the ovary, which is from adult females. Whole ORFeome is plotted as a reference using organismal codon usage frequencies from the Kazusa codon usage database. (**B**) taCAI for human tissues from the Human Protein Atlas. Whole ORFeome is plotted as a reference using organismal codon usage frequencies from the Kazusa codon usage database.

We next examined human genes for tissue-specific differences in codon usage. We obtained transcriptomic data from the Human Protein Atlas (HPA) project and computed taCAI for each tissue. After correcting for the larger number of distinct human tissues with RNAseq data (37 vs 12, see Methods and **Fig 4-Fig Supplement 1**), the testis again emerged as a unique rare codon-enriched outlier tissue, along with the pancreas (**Fig 4B**). These data argue for a conserved and distinctive regulation of mRNAs from rare codon-enriched genes in the testis.

### Testis-restricted expression of the evolutionarily young gene *RpL10Aa* by rare codons impacts fertility

The abundance of rare codon-enriched mRNAs in the testis suggests that rare codons could limit expression of an mRNA/protein to this tissue. Given this, we next searched for an example where testis restriction by rare codons is critical to the animal. The “out-of-testis hypothesis” (Assis & Bachtrog, 2013; Kaessmann, 2010). posits that the testis provides a permissive environment for evolution of new genes. This is based on observations in numerous organisms that young genes tend to be restricted in expression to the testis and gradually gain broader tissue expression patterns as they age (Betrán et al., 2002; Kondo et al., 2017; Marques et al., 2005). The restriction of young genes to the testis has been largely considered a passive consequence of permissive chromatin states during meiosis in the germ cells (Soumillon et al., 2013). However, the reduced impact of rare codons on mRNA and protein expression in the testis may provide an alternative mechanism for restricting the products of young genes to this tissue.

We next explored this possibility that the distinctly higher tolerance for rare codon usage in the testis can restrict expression of evolutionarily young genes. To do so, we analyzed the codon content of young genes arising from duplication events (Zhang et al., 2010). Following a duplication event, there is a parent gene copy that often retains its ancestral function and a child gene copy which may evolve a new function. We analyzed the codon content of evolutionarily young parent and child gene pairs identified by Zhang et. al. (2010), separated by those arising from retrotranspositions versus DNA duplications. We rank-ordered all of the coding sequences in the genome by CAI and plotted the change in genomic CAI rank from parent to child. These data reveal that child genes arising from retrotransposition have lower CAI values than their parent genes (**Fig 5A, DNA duplications vs All Retrotranspositions**). We found that in roughly half (44/96) of these retrotransposition events (**Fig 5A, Retro-ChildMaxInTestis**), child genes have maximum expression in the testis among all tissues analyzed (N=12 tissues), while their parent genes were highest expressed in a tissue other than testis (**Table S3**). Among all retrotransposition-driven duplications, child genes that are maximally expressed among all tissues in the testis are enriched in low CAI values relative to parent genes (**Fig 5A, compare low tails of All Retrotranspositions vs. Retro-ChildMaxInTestis**). These findings imply that rare codons could be functional in restricting expression of evolutionarily young genes to the testis.

**Figure 5.**
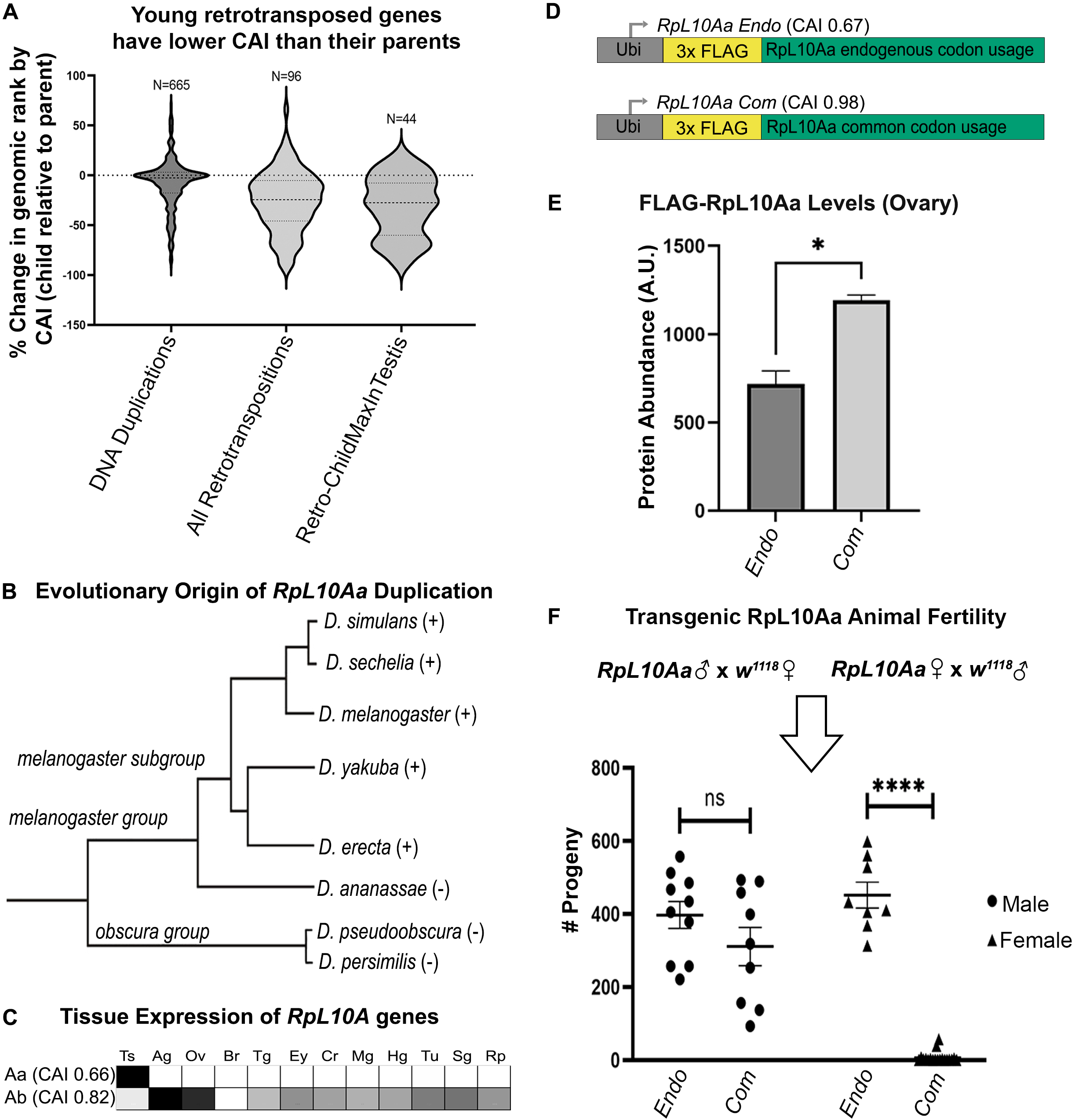
Rare codon usage in evolutionarily young testis genes restricts expression and can impact fertility. (**A**) Codon usage analysis for young parent-child gene duplicate pairs, either arising from DNA duplication (“DNA Duplications”) or from retrotransposition (“All Retrotranspositions”), from Zhang et. Al., 2010. Y axis is percent change in genomic ranking based on difference between child and parent gene CAI values. Direction of change reflects shift in child CAI relative to parent gene. The “Retro-ChildMaxInTestis” dataset is a subset of the “All Retrotranspositions” dataset, and reflects cases where a child gene but not its parent has maximum expression in testis. (**B**) Phylogenetic tree (Carareto et al., 2009) of *RpL10Aa* duplication event. *RpL10Aa* is only present in *melanogaster* subgroup as indicated by (+). *RpL10Aa* is not present in *D. ananassae* or *obscura* group as indicated by (-). (**C**) Expression profiles for *RpL10Aa* and *RpL10Ab* based on FlyAtlas2 RNAseq. Darker boxes indicate higher expression. Ts = testis, Ag = accessory gland, Ov = ovary, Br = brain, Tg = thoracicoabdominal gland, Ey = eye, Cr = crop, Mg = midgut, Hg = hindgut, Tu = Malpighian tubule, Sg = salivary gland, Rp = rectal pad. (**D**) Design of *RpL10Aa* transgenes. CAI for transgenic *3xFLAG-RpL10Aa* CDS indicated in parentheses. (**E**) Quantification of transgenic RpL10Aa protein levels in young (0-8 hours old) adult ovaries measured by western blot. Protein levels normalized to total protein stain. (2 replicates, N=10 animals each, plotting mean +/-SEM, unpaired T-test, p<0.05). Representative blot image in **Figure 5-Figure Supplement 1**. (**F**) *RpL10Aa* fertility assay. Single *RpL10Aa* transgene expressing flies were crossed to wildtype flies of the opposite sex. Adult progeny resulting from a 10-day mating period were counted. Each point represents progeny from one mating pair. (8-19 replicates per condition, Tukey’s, ****p<0.0001). Legend indicates sex of *RpL10Aa* transgene expressing parent.

In many cases, the difference between parent and child can span nearly the entire range of CAI for expressed genes (with the parent at the common end and child at the rare end, **Fig 5A-lower tail of plot for All Retrotranspositions**). From these examples that are present in the Retro-ChildMaxInTestis dataset (**Fig 5A**), we chose a candidate gene to test the idea that codon content restricts expression of a young gene to the testis. The gene encoding the testis-specific ribosomal subunit RpL10Aa has the second largest difference in CAI between parent and child gene of any young retrotransposition mediated duplication in *Drosophila*. The ancestral copy of *RpL10Aa, RpL10A*, is a highly conserved member of the 60s ribosomal subunit. Within the subgenus *Sophophora,* approximately 10 million years ago a gene duplication event, likely mediated by retrotransposition (Marygold et al., 2007; Zhang et al., 2010), gave rise to two isoforms in the *melanogaster* subgroup (**Fig 5B**). This duplication produced a parent gene copy, *RpL10Ab*, and a child copy *RpL10Aa*. We chose to study *RpL10Aa* because of its young evolutionary age, and because of the striking differences in codon usage and tissue expression pattern between *RpL10Aa* and the parent gene *RpL10Ab*. *RpL10Aa* is highly enriched in rare codons (CAI 0.66) placing it in the rarest 8 percent of genes organism-wide, is testis-specific in expression pattern (**Fig 5C**), and is very highly expressed in the testis, ranking in the top 2.5 percent of all genes expressed in the testis. In contrast, *RpL10Ab* has very few rare codons (CAI 0.82), ranking it above 95 percent of genes organism-wide, and is expressed relatively uniformly throughout the body (**Fig 5C**). That such a drastic reduction in codon bias coupled with high testis-specific gene expression occurred over a short evolutionary period suggests that rare codons may have been positively selected for in *RpL10Aa* for the purpose of restricting protein levels outside of the testis.

To test the hypothesis that rare codon enrichment in a young gene can drive testis-specificity, we analyzed protein levels for both an endogenous *RpL10Aa* sequence (*RpL10Aa Endo*) and a codon optimized *RpL10Aa* sequence with all common codons (*RpL10Aa Com*). We did this by placing both coding sequences in the same *ubi*-promoter used for our organism-wide GFP reporter screen (**Fig 5D**). To specifically measure RpL10Aa protein from transgenes and not from the endogenous gene locus, we included an N-terminal 3x-FLAG tag in both the endogenous and codon optimized *RpL10Aa* sequences. While generating stable genomic lines of the *RpL10Aa* reporters, we noticed that *RpL10Aa Com* animals are much less fertile than *RpL10Aa Endo* animals. This led us to measure protein levels of both reporters in the ovary by western blot. We find that RpL10Aa Com protein is 2-fold higher than RpL10Aa Endo protein in ovaries (**Fig 5E, Fig 5-Fig Supplement 1**). This finding is consistent with the idea that endogenous rare codon content is sufficient to limit protein levels in tissues outside of the testis.

Our previous study of codon-dependent effects of Ras signaling in *Drosophila* identified profound phenotypic differences of a 2-fold, codon-dependent increase in protein expression (Sawyer et al., 2020). We next examined the impact of increased, codon-dependent protein expression of RpL10Aa. Taken together with our initial observation that *RpL10Aa Com* animals are less fertile than *RpL10Aa Endo* animals, we hypothesized that female fertility but not male fertility is impacted by codon optimization of the *RpL10Aa* coding sequence. To evaluate male and female fertility for *RpL10Aa Com* and *RpL10Aa Endo* animals, we performed reciprocal single fly crosses between our transgenic flies and wildtype flies. Throughout the course of our fertility assay, we observed high female lethality in *RpL10Aa Com* animals (11/30 females, 1/10 males) and low lethality in *RpL10Aa Endo* animals (2/10 females, 0/10 males). Only crosses where both parents survived the duration of the study were analyzed with respect to fertility. We observe no difference in male fertility between endogenous and codon optimized *RpL10Aa* animals when crossed to wildtype females (**Fig 5F**). For the reciprocal cross (wildtype males crossed to transgenic females) we instead observe drastic differences in female fertility between reporters. *RpL10Aa Com* females produced just 6 viable offspring on average compared with *RpL10Aa Endo* females, which produced on average 452 viable offspring (**Fig 5F**). Therefore, the endogenously encoded rare codon content is capable of restricting RpL10Aa expression in tissues outside of the testis. Altering codon content of this testis-specific gene causes female infertility (see **Discussion**). Taken together, our findings illustrate an important role for rare codons in regulating the tissue-specificity of protein expression and imply that rare codons play a particularly important role in testis biology.

## Discussion

Redundancy of the genetic code has long been a mystery of the central dogma of molecular biology. Here, we use *Drosophila melanogaster* to uncover distinct tissue-specific responses to the same genetic code. Our work uncovers fundamental aspects of rare codon biology. We reveal a cliff-like limit on rare codon usage per gene. Near the boundaries of this limit, however, we demonstrate that individual tissues have different tolerances for protein expression from genes enriched in rare codons. Tissues with a high tolerance for rare codons include the brain of both sexes and the testis, but not the ovary. Taking a closer look at the testis, we find the male germ cells and somatic cells of the germline stem cell niche of the testis robustly express protein from rare codon-enriched reporters, while the somatic cyst cells do not. In developing a new metric for tissue-specific codon usage, taCAI, we reveal endogenous genes in the testis of both *Drosophila* and humans show an abundance of rare codon-enriched mRNAs, suggesting that rare codon-enriched genes play an essential and conserved role in testis biology. In search of a physiologically relevant role for rare codon tolerance in testis biology, we demonstrate that endogenously rare codons can restrict the expression of the evolutionarily young gene *RpL10Aa* to the testis. This restriction proves to be crucial for female fertility and organism viability. Taken together, our results uncover clear tissue-specific differences in the impact of codon usage bias and support a novel role for rare codons in restricting the expression of evolutionarily young genes to the testis.

### Codon bias as an important parameter for organism-wide and tissue-specific mRNA and protein expression

Here, our sensitive reporter-based approach suggests a strong correlation between CAI and reporter GFP abundance in whole *Drosophila* (r^2^=0.82, p<.0001). We note that previous data from bioinformatic approaches in human tissues are conflicting regarding correlations between CAI and tissue-specific mRNA and protein expression (Plotkin et al., 2004; Sémon et al., 2006). Further, similar bioinformatic studies suggested only weak codon-dependent differences between tissues in *Drosophila* (Payne & Alvarez-Ponce, 2019). The sensitivity of our reporter and the ability of our expression platform to rapidly test various sequences over a narrow CAI range proved helpful in defining rare codon limits for protein expression, which closely mirror existing mRNA expression data for the testis. Our study reveals a strict limit to rare codon usage on a genomic scale. This limit could reflect the tipping point in rare codon usage before competition between transcription and mRNA decay favors degradation over translation (Buschauer et al., 2020; Radhakrishnan et al., 2016).

We also observe clear differences in the impact of codon usage bias between *Drosophila* tissues. Importantly, we show how different tissues respond differently to the same genetic code. It is well appreciated that tissue-specific chromatin environments play critical roles in transcriptional regulation. Our findings here suggest that tissue-specific tolerances for rare codons play a similar role at the level of translation. There are several likely candidate mechanisms underlying tissue-specific impacts of codon bias, including differences in levels of RNA decay machinery or tRNAs between tissues. Several RNA decay pathways have been linked to rare codon usage and would be of interest to follow up on in our system (Buschauer et al., 2020; Radhakrishnan et al., 2016). Levels of expressed tRNAs have been previously measured in *Drosophila* at the organismal level (Moriyama & Powell, 1997; White et al., 1973), but not at the level of individual tissues. Quantifying tRNA expression is not the same as measuring tRNA activity, however, as tRNAs exhibit rigid folding structure, distinct amino acid carrying status when active, and require post transcriptional modifications. The contribution of tRNA regulation to tissue-specific codon bias regulation can be explored further in the future.

We note that mechanisms which lead to rare codon mRNA/protein accumulation may be distinct in different tissues. For example, our taCAI analysis of RNAseq data revealed that the testis but not the brain is enriched for mRNAs with abundant rare codons. Recent work by other groups, however, demonstrates that rare codons have less influence on mRNA stability in central nervous system compared to whole embryo (Burow et al., 2018), supporting our finding here that the brain has unique biology pertaining to rare codons. Further, while the accessory gland did not appear to express our rare codon-enriched reporters, our taCAI analysis suggests this male reproductive tissue is also enriched in mRNAs with abundant rare codons. Now that we have pinpointed interesting differences between the male reproductive tract, brain, and other tissues, these differences can be exploited in future studies to reveal the underlying molecular mechanisms.

### Rare codons and gene evolution-a potential role for the testis

Here, we find a conserved enrichment for rare codons in testis-specific genes. Further, using the evolutionarily young gene *RpL10Aa* as an example, we demonstrate a role for rare codons in restricting gene expression to the testis. Optimizing the codon sequence of this testis-specific gene causes ectopic high expression in the ovary and results in female infertility. We speculate that ectopic RpL10Aa in the ovary may act as a dominant negative relative to the function of the parent gene *RpL10Ab* (64% identity between these proteins), which is required for female fertility (Wonglapsuwan et al., 2011). This role for tissue restriction of expression by rare codons may have functional relevance to the “out-of-testis” hypothesis for new gene evolution (Kaessmann, 2010). This hypothesis proposes that the testis acts as a “gene nursery” to promote evolution and neofunctionalization of young genes through its permissive expression environment and intense selective pressures from both mate and sperm competition. This is based on observations that testes are the fastest evolving tissue and that young genes are often restricted in expression to the testis and gain broader expression patterns as they age.

While the current model is that permissive chromatin states passively allow expression of young genes resulting from retrotransposition events specifically in the testis (Kaessmann, 2010), here we demonstrate that rare codon content can also fulfill this role. We suggest that rare codons could be positively selected for in evolutionarily young genes in *Drosophila*. Male-biased genes have also been characterized to have rarer codon usage than female-biased genes and non-sex-biased genes in numerous plant species (Darolti et al., 2018; Song et al., 2017; Whittle et al., 2007) and *Drosophila* (Hambuch & Parsch, 2005), and have rarer codon usage than non-sex-biased genes in zebrafish and stickleback (Yang et al., 2016). Thus, our findings may be widely applicable to sexually reproducing species across phylogenetic kingdoms. Taken together, we highlight codon usage as a critical and under-appreciated aspect of tissue-specific gene regulation that merits greater attention, notably in the fields of evolutionary and developmental biology.

## Materials and Methods

### Generation of codon-modified reporters in Drosophila

Codon-modified exon sequences for *GFP* and *RpL10Aa* were designed according to the codon usage frequencies in the *Drosophila melanogaster* genome taken from the Kazusa codon usage database (https://www.kazusa.or.jp/codon/). The *mCherry* sequence used in generating fusion proteins was obtained from NovoPro pADH77. Sequences were subsequently generated through gene synthesis (ThermoFisher Scientific and Invitrogen, Waltham, MA) and cloned into a pBID-Ubi plasmid (modified from Addgene Plasmid #35200) or were directly synthesized into the desired pBID-Ubi plasmid cloning site (Twist Bioscience, South San Francisco, CA). Plasmids were grown in NEB 5-alpha competent *E. coli* cells (NEB #C2987), transformed according to the manufacturer’s protocol, and purified with a ZymoPure II Plasmid Midipred Kit (Zymo Research). Plasmids were injected into *attP40 (2L)* flies by Model System Injections (Durham, NC). Full sequences for the pBID-Ubi plasmid and synthesized genes are provided in **Table S1**.

### Fly stocks

All flies were raised at 25°C on standard media unless noted otherwise (Archon Scientific, Durham, NC). The following stock was obtained from the Bloomington Drosophila Resource Center *w^1118^* (#3605). The following stocks were generated for this study: *GFP30D*, *GFP60Dv1*, *GFP60Dv2*, *GFP60Dv3*, *GFP70D*, *GFP80D*, *GFP90D*, *GFP100D*, *GFP50C5’*, *GFP50C3’*, *GFP60C3’*, *GFP70C3’*, *GFP80C3’*, *GFP90C3’*, *GFP54C3’*, *mGFP100Dv1*, *mGFP100Dv2*, *mGFP100Dv3*, *mGFP100Dv4*, *mGFP100Dv5*, *mGFP100Dv6*, *mGFP100Dv7*, *RpL10Aa Endo*, *RpL10Aa Com*.

### Protein quantifications

Protein samples were prepared as in Sawyer et al., 2020. Briefly, tissues or whole animals were homogenized in Laemmli buffer (10 tissues in 50ul or 5 animals in 100ul) on ice, then boiled for 5 min. Samples were separated on 12% sodium dodecyl sulfate polyacrylamide gels by electrophoresis (SDS-PAGE) at 200V. Proteins were transferred onto nitrocellulose membranes using an iBlot 2 Dry Blotting System (Invitrogen, Waltham, MA) set to 20V, 6min. Total protein was quantified using Revert700 Total Protein Stain Kit (LI-COR Biosciences, Lincoln, NE) according to the manufacturer’s protocol for single channel imaging. For normalization we note that total protein stain was superior to antibodies against single housekeeping genes (Tubulin) due to variable expression of these proteins between tissues. The following antibodies were used: rabbit anti-GFP (1:1,000, Invitrogen, cat#A11122), anti-FLAG M2 (1:500, Sigma, St. Louis, MO, cat#F1804), IRDye 800CW (1:10,000, LI-COR Biosciences, anti-rabbit or anti-mouse). Signal was detected using a LI-COR Odyssey CLx and analyzed using Image Studio version 5.2 (LI-COR Biosciences).

### CAI

CAI calculations for transgenic reporters and endogenous *Drosophila* genes were performed using CAIcal (Puigbò et al., 2008). For CAI calculation of endogenous genes, the standalone version of CAIcal was used with input CDS sequences from the *Drosophila melanogaster* genome version r 6.22, and the *Drosophila melanogaster* codon usage table from the Kazusa codon usage database (Nakamura et al., 2000).

### taCAI

Tissue-specific RNA sequencing data was obtained for *Drosophila* from FlyAtlas2 and for humans from the Human Protein Atlas. We defined genes as being tissue-specific in *Drosophila* if they were present in the 40% highest expressed genes in the tissue of interest and excluded from the 25% highest expressed genes of all other tissues analyzed. Applying this same filter to human tissues revealed that testis genes were most enriched in rare codons among the 37 tissues analyzed (**Fig 4-Fig Supplement 1**), however, the three-fold larger number of tissues in humans compared to flies (37 vs 12) prompted us to explore additional, more stringent cutoffs of tissue specificity for the human data. Towards this end we defined genes as being tissue-specific in human if they were highest expressed in the tissue of interest and had a TPM <1 in all other tissues analyzed (**Fig 4B**). Codon usage frequency was calculated using the Kazusa Countcodon program (https://www.kazusa.or.jp/codon/countcodon.html). We used the resulting tissue-specific codon frequency table to compute taCAI values for each tissue using the following equation:

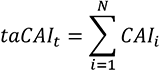

where *N* is the number of genes in the genome and *CAI_i_* is the codon adaptation index value calculated for gene *i* using the tissue-specific codon frequency table for tissue *t*. This calculation returns a genome-sized distribution of values, where the codon usage of each gene is evaluated relative to the codon usage of a pre-defined tissue-specific gene set. The distribution of taCAI values for a given tissue can be visualized as a violin plot and these distributions can be compared between tissues. To additionally visualize how the codon usage of each tissue compares to the codon usage of the entire genome, we computed the genomic taCAI distribution using the following equation:

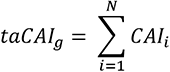

where *N* is the number of genes in the genome and *CAI_i_* is the codon adaptation index value calculated for gene *i* using the codon frequency table for the entire genome *g*.

### qRT-PCR

Animals heterozygous for both m*GFP100Dv1* and *GFP0D* were aged 3-7 days at 25°C on standard fly medium supplemented with wet yeast (Archon Scientific, Durham, NC). Dissections were performed in RNase-Free phosphate buffered saline and completed in under 2 hours. After dissection, tissues were immediately homogenized in TRIzol reagent (N=10 animals per tissue per replicate, 500ul TRIzol reagent) and snap frozen in liquid nitrogen before storage at -80°C. RNA was purified according to the manufacturer’s protocol, using glycogen as a carrier and resuspending in molecular grade water. RNA was then treated with DNase I at room temp for 15 minutes before terminating the reaction by adding 2.5mM EDTA and incubating at 65°C for 10 mins then storing at -80°C. Quantification of RNA was performed on a NanoDrop spectrophotometer and samples were diluted to match the concentration of the lowest concentration sample. Equal amounts of RNA for all samples in **Fig 3N** were simultaneously transcribed into cDNA using iScript cDNA synthesis kit (BIO-RAD, Hercules, CA, cat#170-8891) according to the manufacturer’s protocol within 7 days of the initial dissections to preserve sample quality. No Reverse Transcriptase (NRT) controls were also run simultaneously for each sample to control for genomic DNA contamination. Quantitative Real-Time PCR (qRT-PCR) was run simultaneously on all samples in **Fig 3N**, corresponding NRT controls, and No Template Controls (NTC) for each primer pair using Luna Universal qCPR Master Mix (NEB, Ipswich, MA, #M3003) following the manufacturer’s protocol (1ul cDNA, 36.5ng, per 10ul reaction). A CFX384 Touch Real-Time PCR Detection System (BIO-RAD) was used for cDNA amplification and detection of FAM/SYBR Green fluorescence. Primers were designed against the GFP segment of m*GFP100Dv1* and against *GFP0D*. *mGFP100Dv1* qPCR FW primer: AGGGGAAGAATTATTTACTGGGGT, *mGFP100Dv1* qPCR RV primer: CCCATAAGTTGCGTCCCCTT, *GFP0D* qPCR FW primer: GGCAAGCTGACCCTGAAGTT, *GFP0D* qPCR RV primer: TTCATGTGATCGGGGTAGCG. qRT-PCR run data was analyzed using BIO-RAD CFX Manager software. Amplification curves for all samples had characteristic S-shapes and each qPCR primer pair yielded a single melting peak. All cDNA samples reached threshold several cycles earlier than NRT and NTC controls. Technical replicates for all samples yielded very consistent CT values. A single testis sample, however, yielded an outlier 2^(-delta delta CT) value (Tukey’s p<.0001), and was removed from the analysis after bleach-agarose electrophoresis revealed that this sample lacked detectable RNA. Relative transcript abundance was calculated using the 2^(-delta delta CT) method. Graphs were generated using GraphPad Prism 9.2.0.

### Fluorescence Imaging

Whole larval images were acquired on either a Zeiss SteREO Discovery.V12 (Zeiss Achromat S 0.63x FWD 107mm objective) and Zeiss AxioCam ICc 5 camera (images in **Fig 1**), or on a Leica MZ10 F stereoscope (Leica Plan APO 1.0x objective #10450028) and Zeiss AxioCam MRc r2.1 camera (images in **Fig 2**). To immobilize larvae for imaging, wandering L3 larvae were anesthetized using di-ethyl ether for 3.5 minutes as described in (Kakanj et al., 2020). Larval and adult tissues were prepared for imaging by dissecting in 1x PBS, fixing in 1x PBS, 3.7% paraformaldehyde, 0.3% Triton-X for 30 minutes, and staining with Hoechst 33342 (1:1,500, Life Technologies, Carlsbad, CA, #c10339). Tissues were mounted in Vectashield (Vector Laboratories Inc., Burlingame, CA). Tissue images were acquired on an upright Zeiss AxioImager M.2 without Apotome processing and with the Apotome unit positioned out of the light path (Zeiss 10x NA 0.3 EC Plan-Neofluar objective) using a Zeiss Axiocam 503 mono version 1.1. Imaging conditions were identical between tissues to be visually compared (noted in the figure legends). Image processing was performed in FIJI (formatting) and Photoshop (applying levels adjustment layer uniformly across panels to be visually compared).

## Key Resources Table

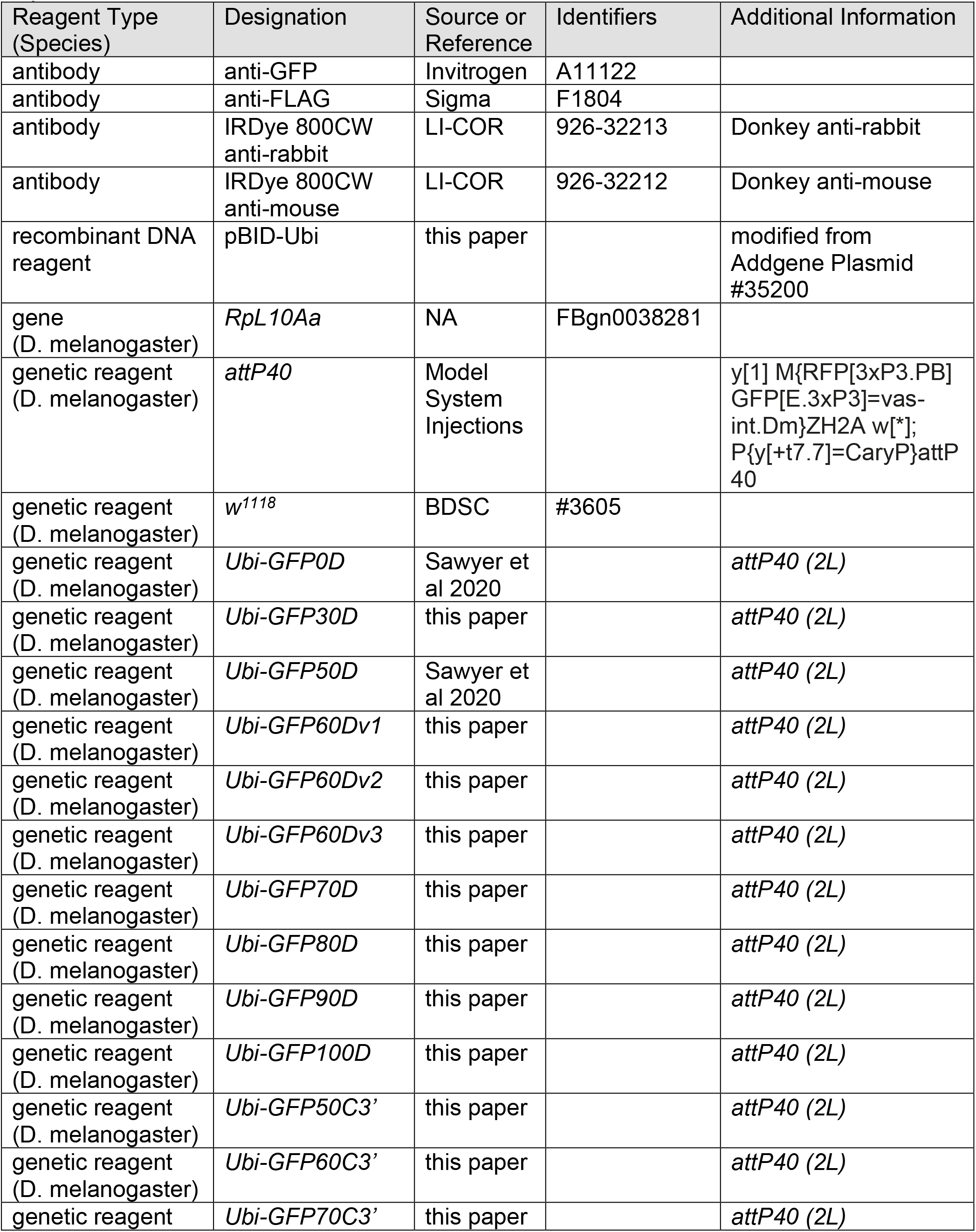
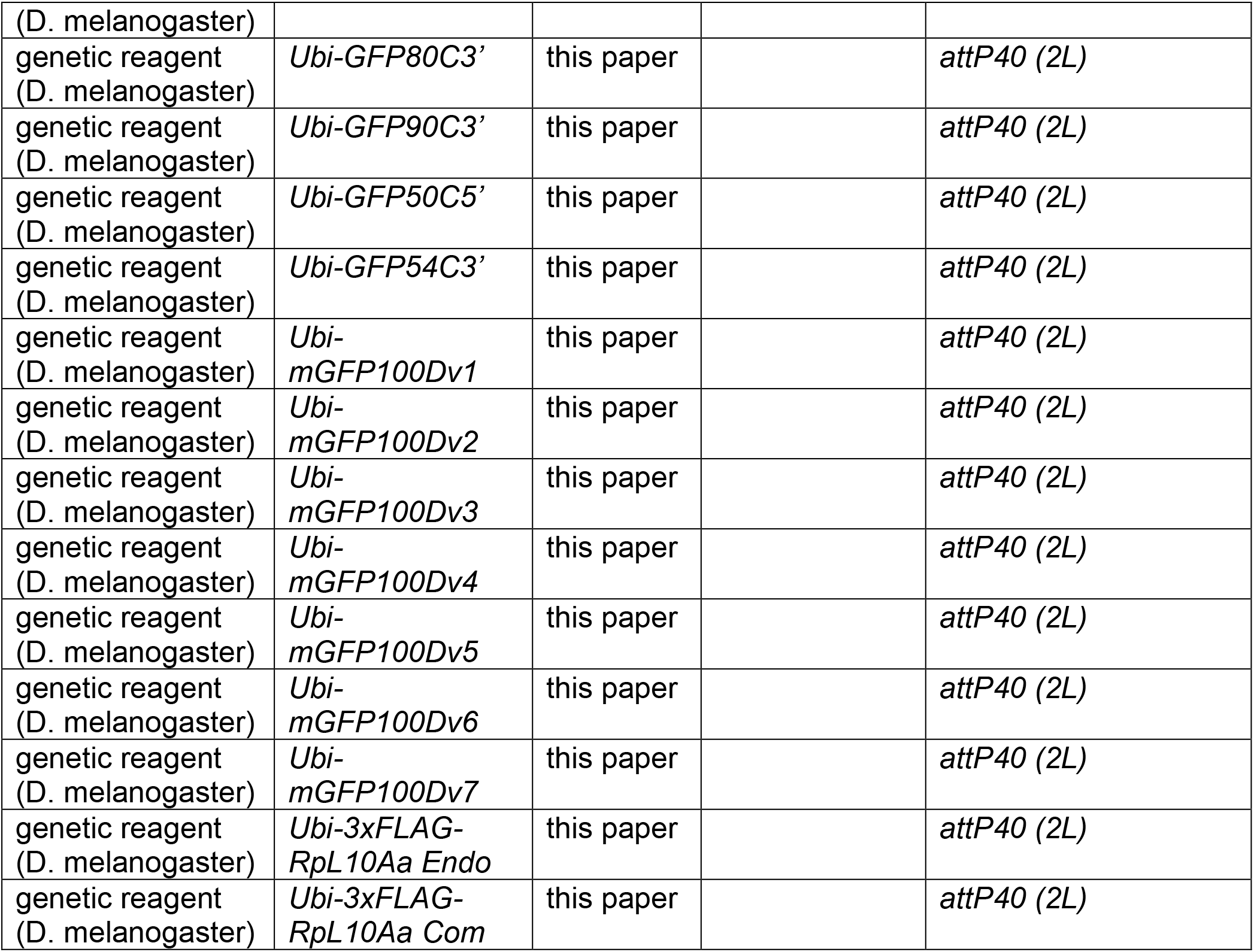

## Acknowledgments

The following kindly provided reagents used in this study: Bloomington Drosophila Stock Center, Developmental Studies Hybridoma Bank. David MacAlpine (Duke) provided valuable technical guidance for RT-PCR. Braden Tierney (Cornell) provided technical support in downloading FlyAtlas2 data. We thank Danny Lew and Zhao Zhang (Duke) for comments on the manuscript. This project was supported by ACS grant RSG-128945 to D.F., an NSF GRFP grant to S.A., NIH grants R01CA94184 and P01CA203657 to C.M.C., and NIHG grants R35 GM140844 and R01 HL111527 to A.L.

## Competing Interests

The authors declare no competing interests.

## Supplemental Data

**Figure 1 – Fig Supplement 1.**
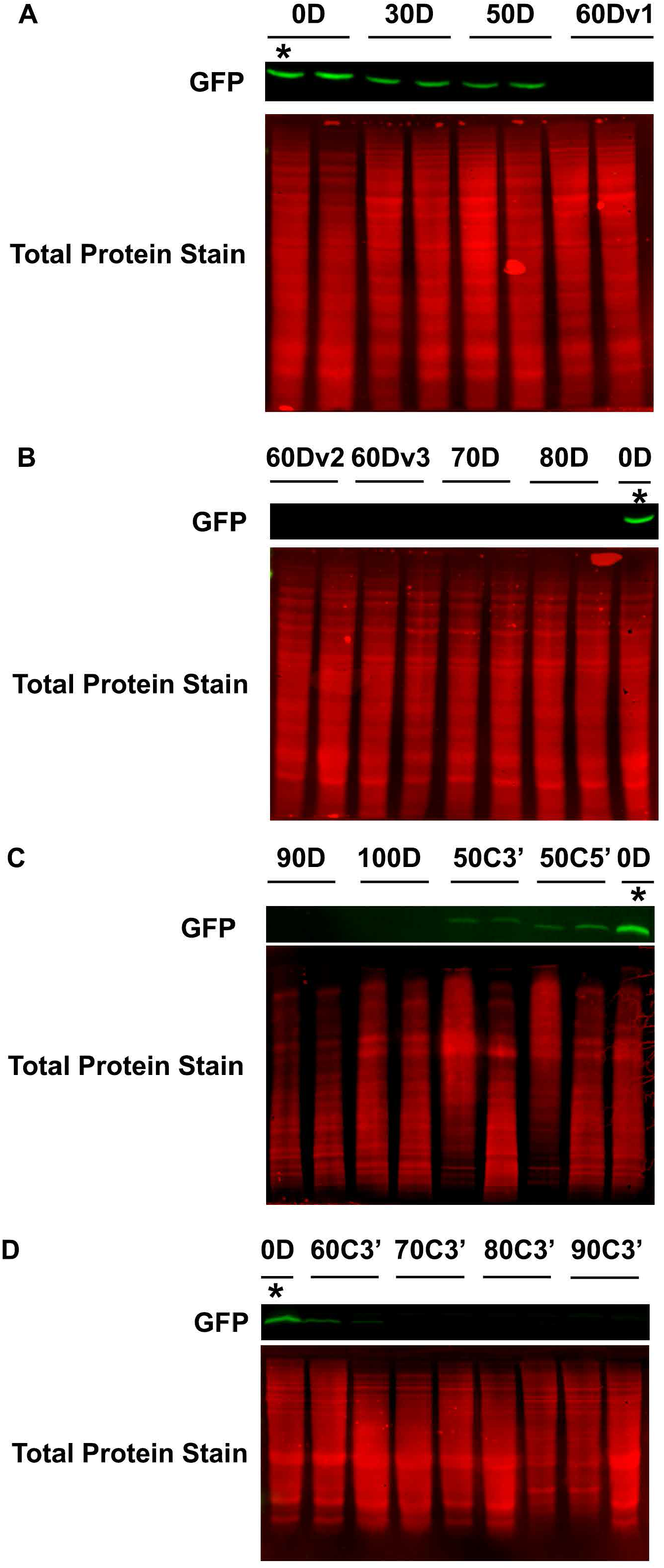
Western blots of whole WL3 larva. (**A-D**) Western blots of whole larvae with stable genomic insertions of the indicated reporters, performed in duplicate. Detected GFP signal is indicated by green bands. Total protein stain for each lane is displayed in red directly below the GFP signal bands. *indicates the same GFP0D lysate loaded in each gel for between gels comparisons.

**Figure 2 – Fig Supplement 1.**
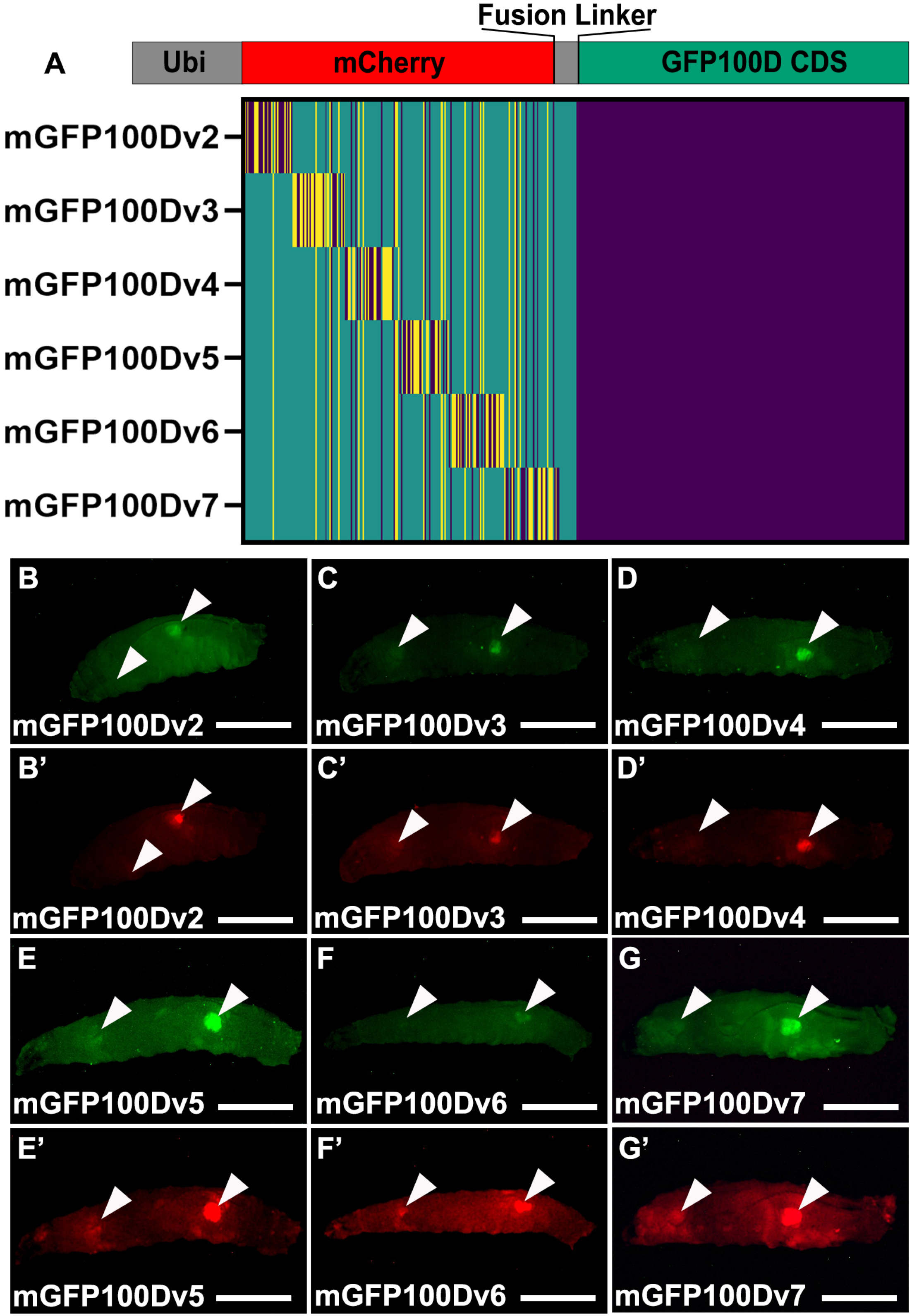
Tissue specific rescue of GFP100D is not dependent on mCherry or linker sequence. (**A**) Schematic of each indicated reporter. Heat map color scheme as in **Fig 2A**. (**B-G**) Fluorescent images taken in the GFP excitation channel of live male WL3 larva with stable genomic insertions of the indicated reporter. (**B’-G’**) Fluorescent images taken in the mCherry excitation channel of live male WL3 larva with stable genomic insertions of the indicated reporter. Arrowheads indicated tissues that robustly produce the fluorescent reporter compared.

**Figure 3 – Fig Supplement 1.**
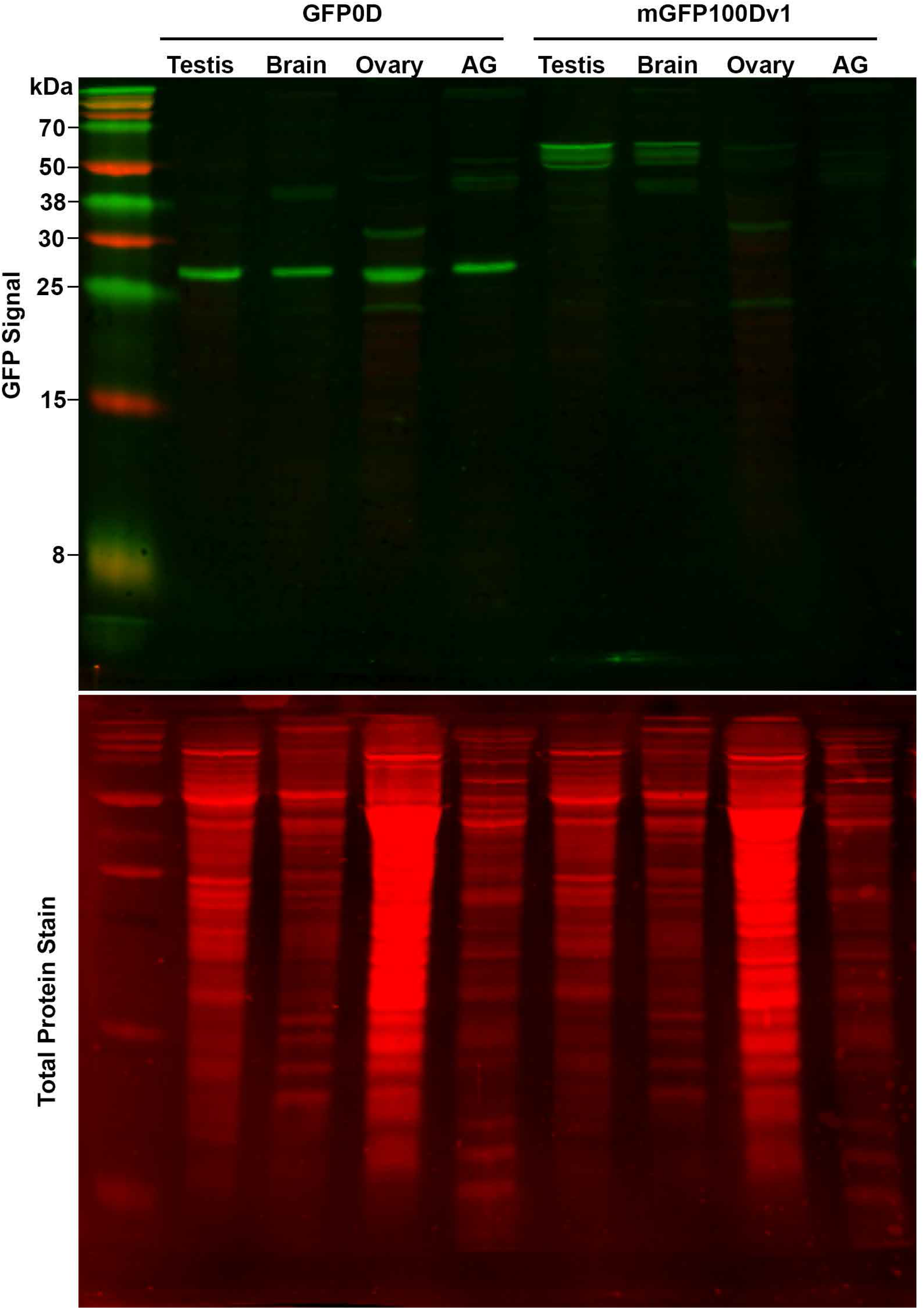
Western blot for adult tissues. Representative western blot for dissected adult tissues (N=10-12 per tissue, 10-12 pairs for testis, ovary, and AG, three replicates done in total, quantified in **Fig 3M**). GFP signal is indicated by green bands. For GFP0D the prominent band around 27 kDa was quantified. For mGFP100Dv1, the three closely adjacent bands around 55 kDa were quantified. Total protein stain for each lane is displayed in red directly below the GFP channel image.

**Figure 3 – Fig Supplement 2.**
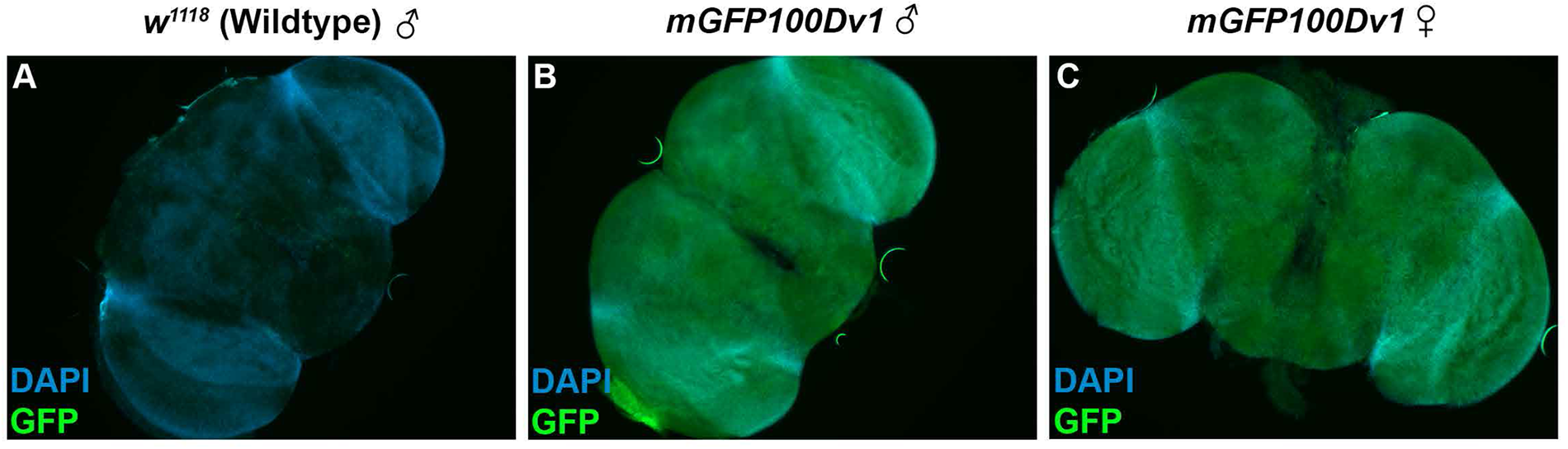
The brains of both sexes robustly express rare codon-enriched reporters. (**A**) Fluorescent image of a dissected brain from an adult *Drosophila* with no transgenic GFP reporter. Image is taken under identical conditions as **B-C** with fluorescence intensity normalized to male *mGFP100Dv1* brain. Nuclei are stained with DAPI, indicated in blue. GFP channel signal, indicated in green, represents autofluorescence. (**B-C**) Fluorescent images of dissected brains from adult *Drosophila* with stable genomic insertion of *mGFP100Dv1*. Images are taken under identical conditions with fluorescence intensity normalized to male *mGFP100Dv1* brain. Nuclei are stained with DAPI, indicated in blue. GFP channel signal, indicated in green, is from the *mGFP100Dv1* reporter.

**Figure 4 – Fig Supplement 1.**
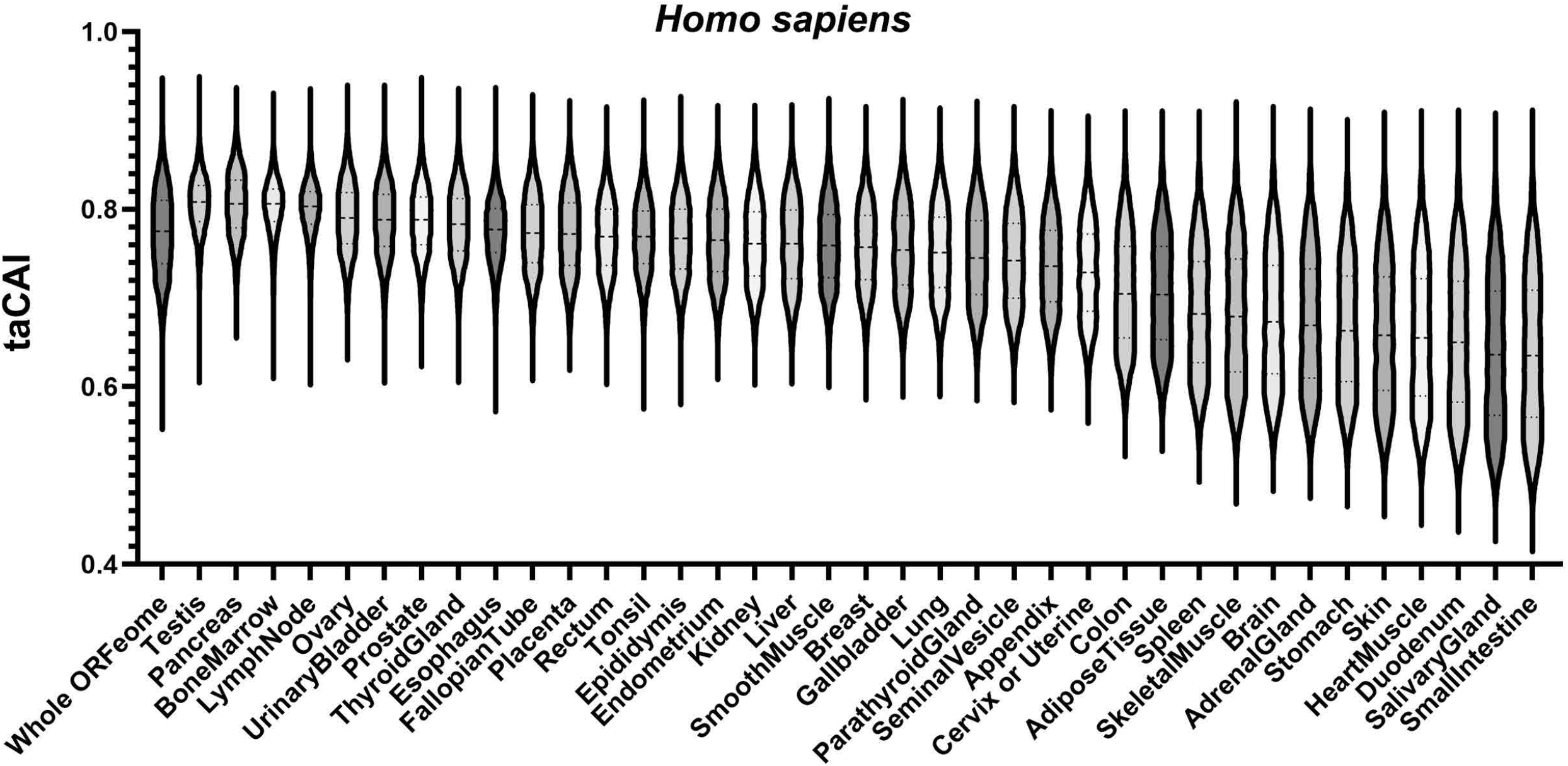
Human taCAI using identical cutoffs as fly. taCAI for human tissues from the Human Protein Atlas. Whole ORFeome is plotted as a reference using organismal codon usage frequencies from the Kazusa codon usage database. The criteria for determining tissue-specific gene sets here are identical to the criteria for determining tissue-specific gene sets in *Drosophila melanogaster* (see Methods).

**Figure 5 – Fig Supplement 1.**
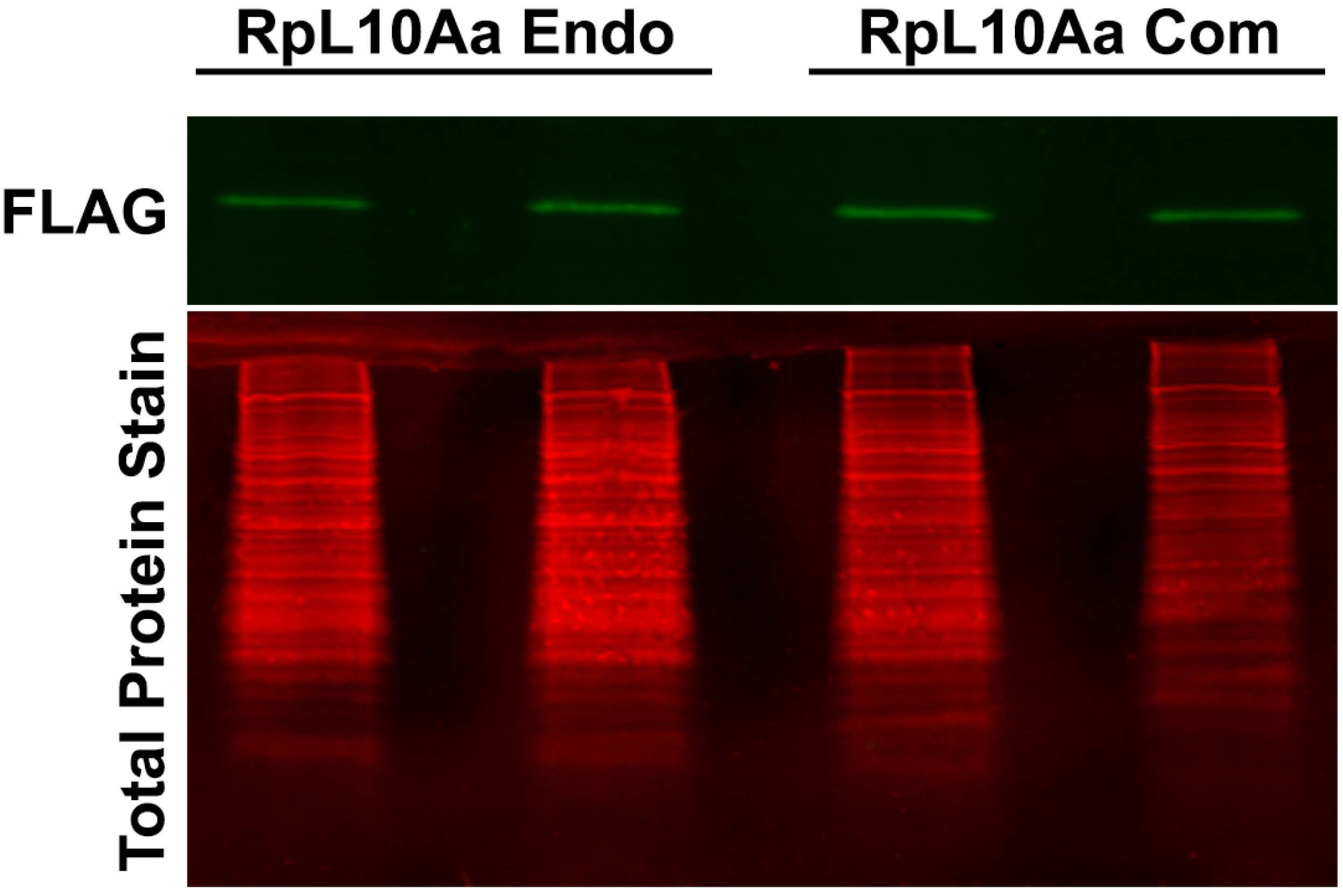
Western blot for RpL10Aa in adult ovaries. Western blot of adult ovaries from animals with stable genomic insertion of *RpL10Aa Com* and *RpL10Aa Endo* (2 replicates, N=10 pairs of ovaries each, quantified in **Fig 5E**)

**Table S1.**
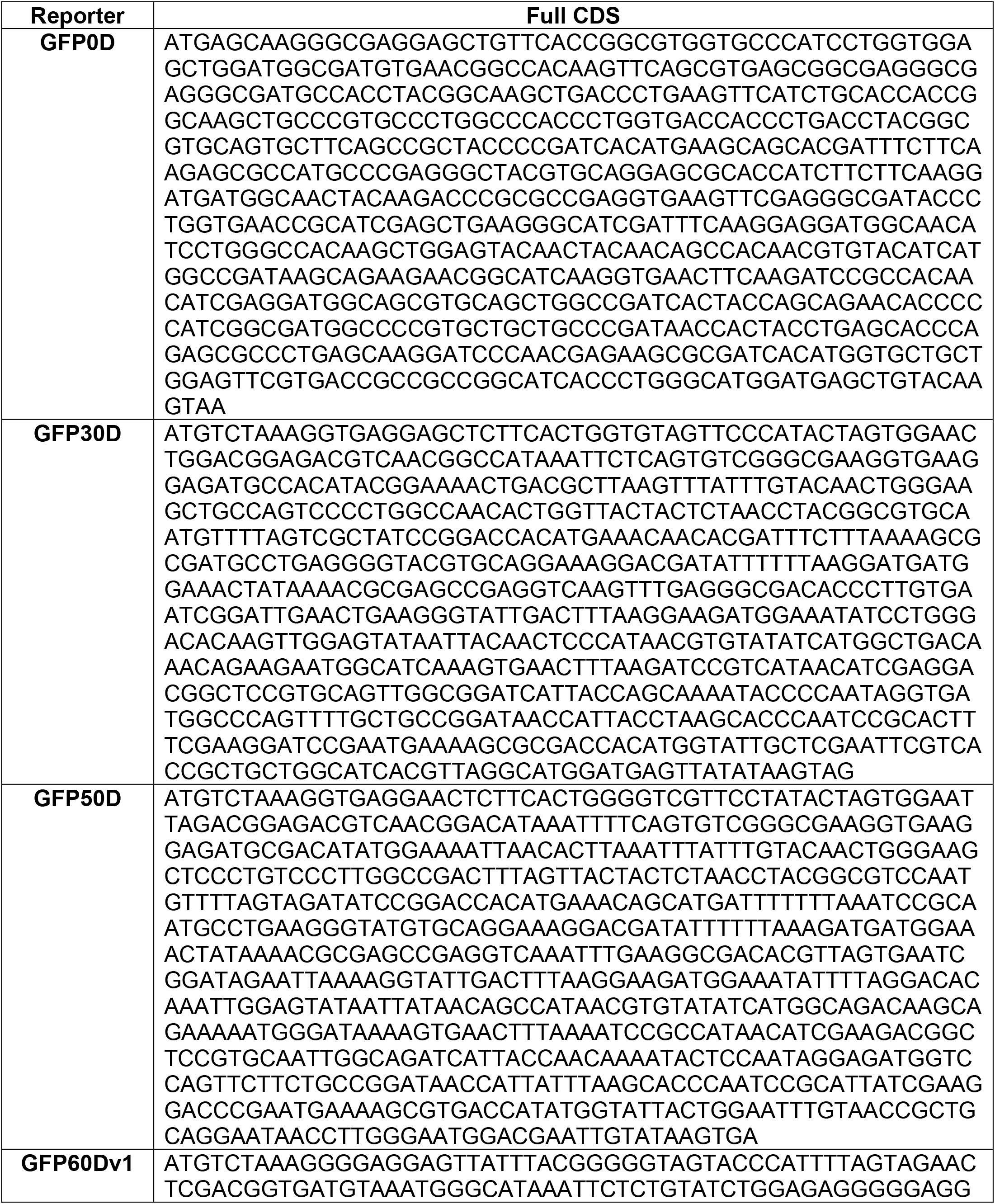
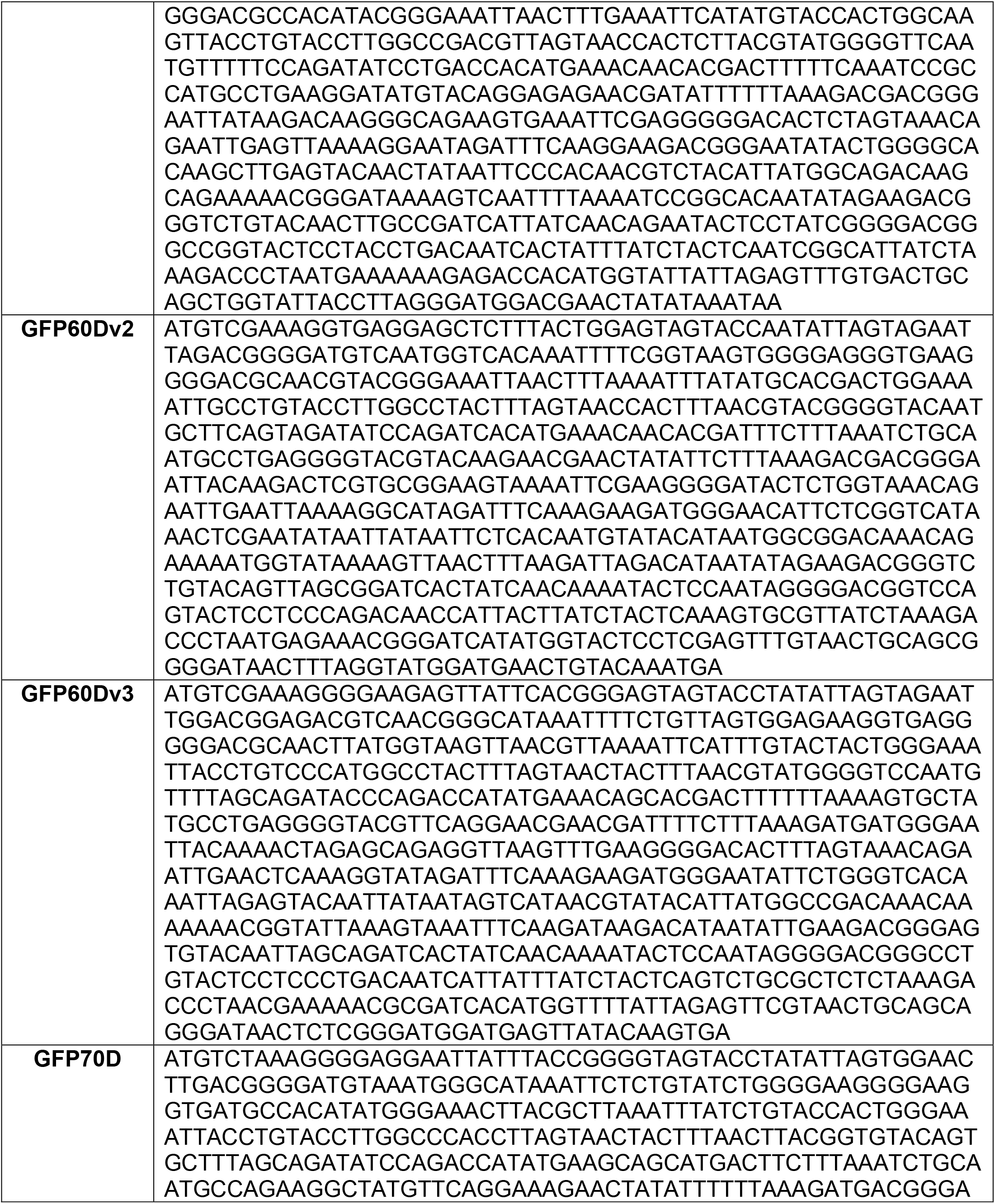
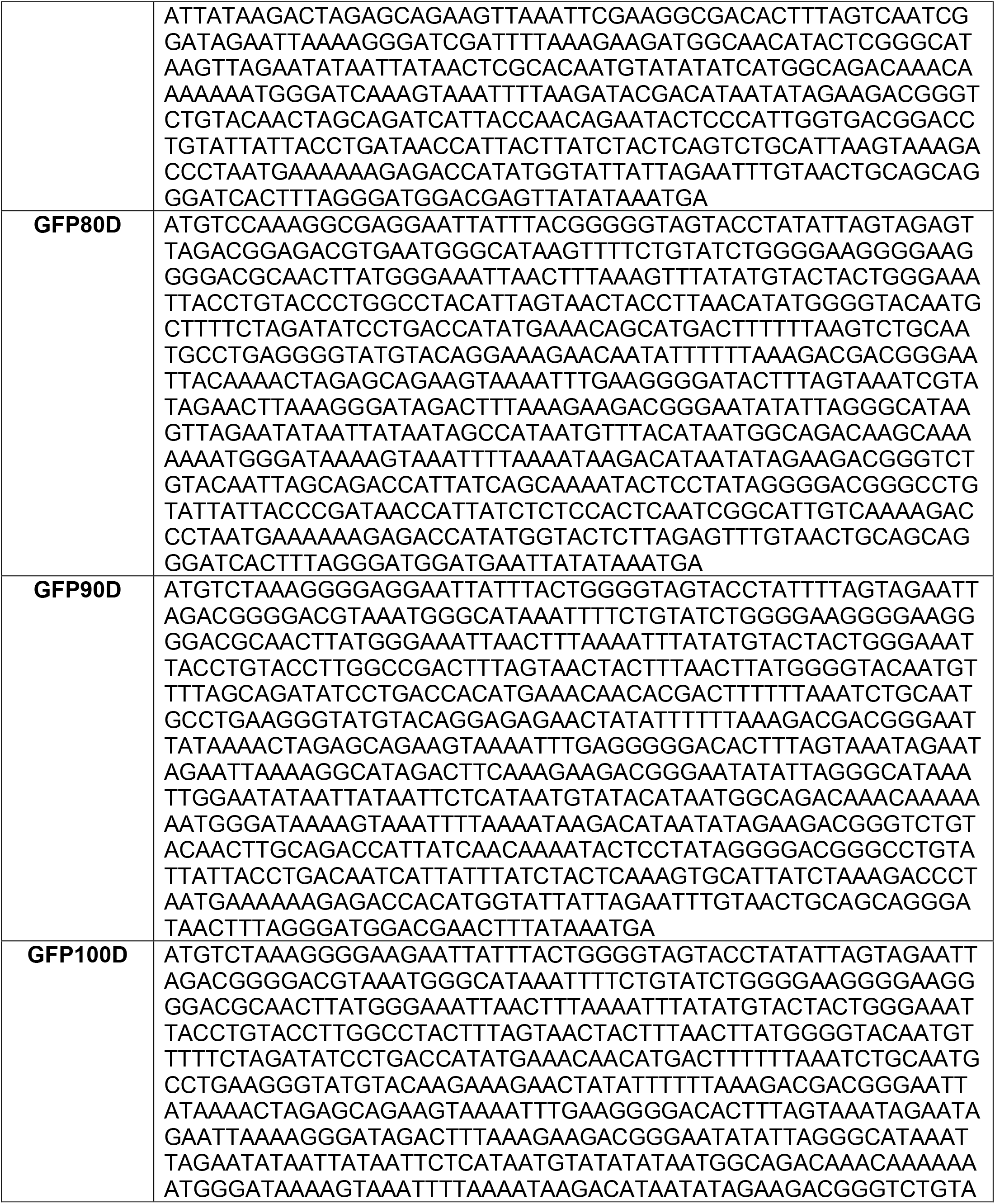
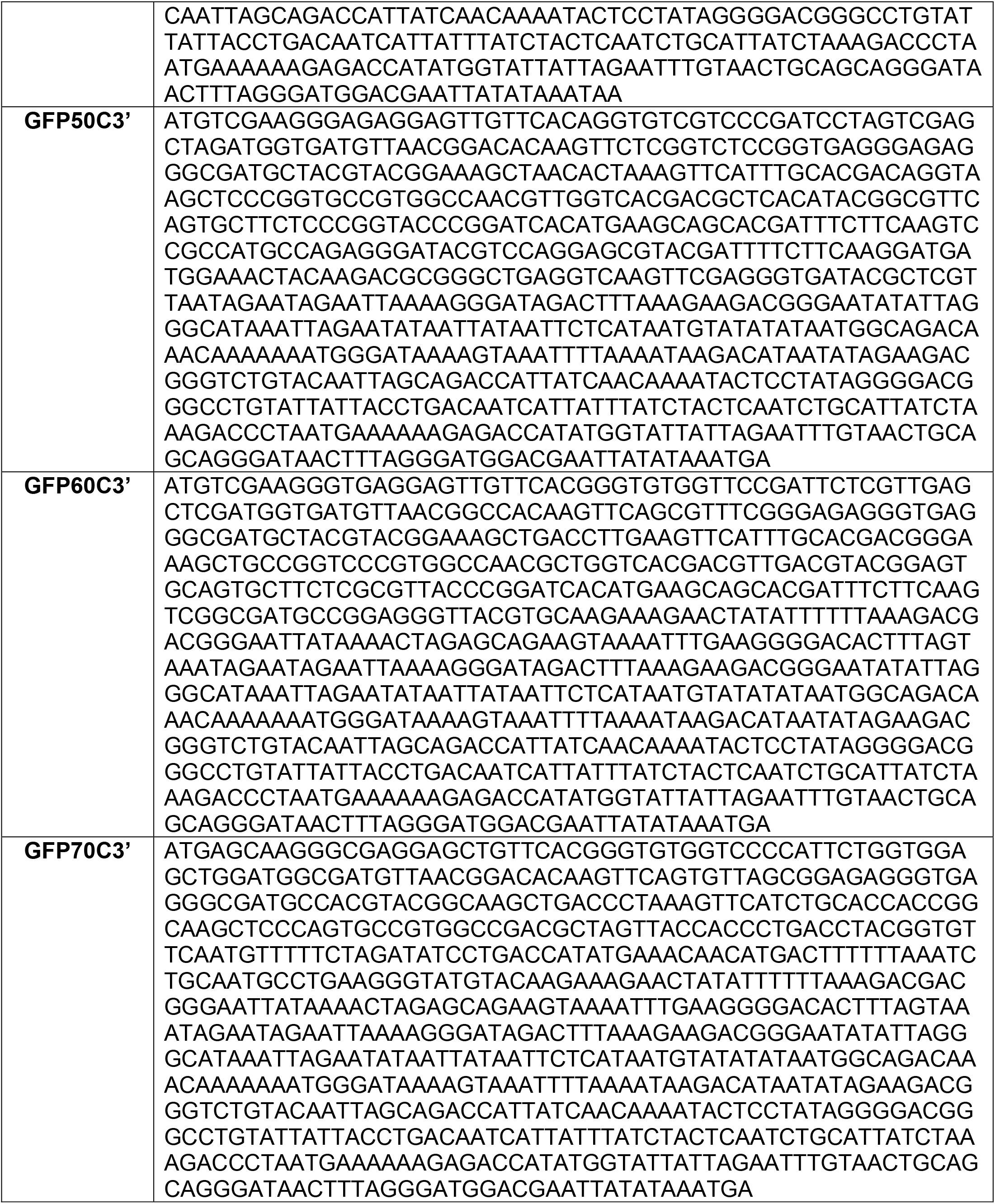
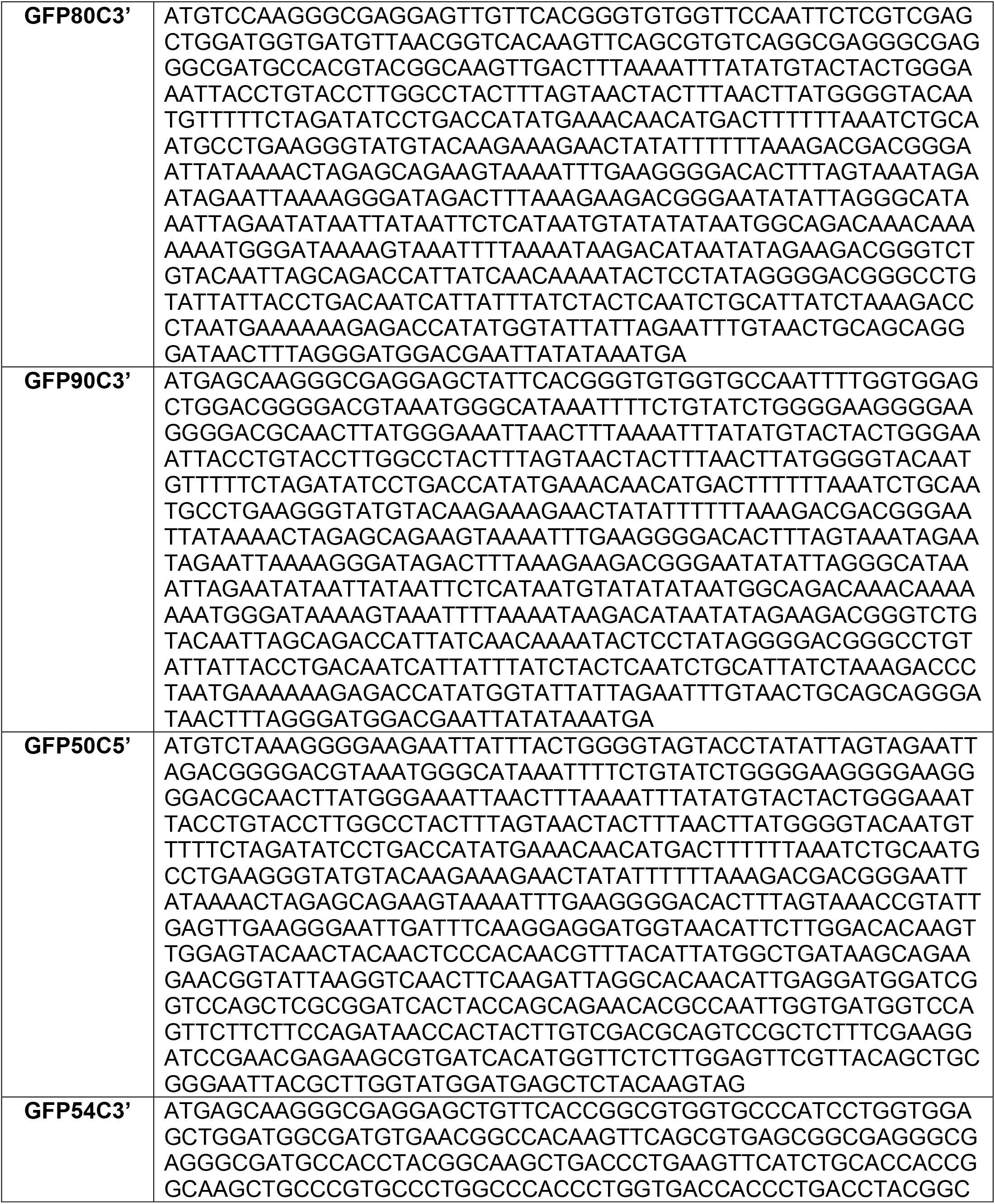
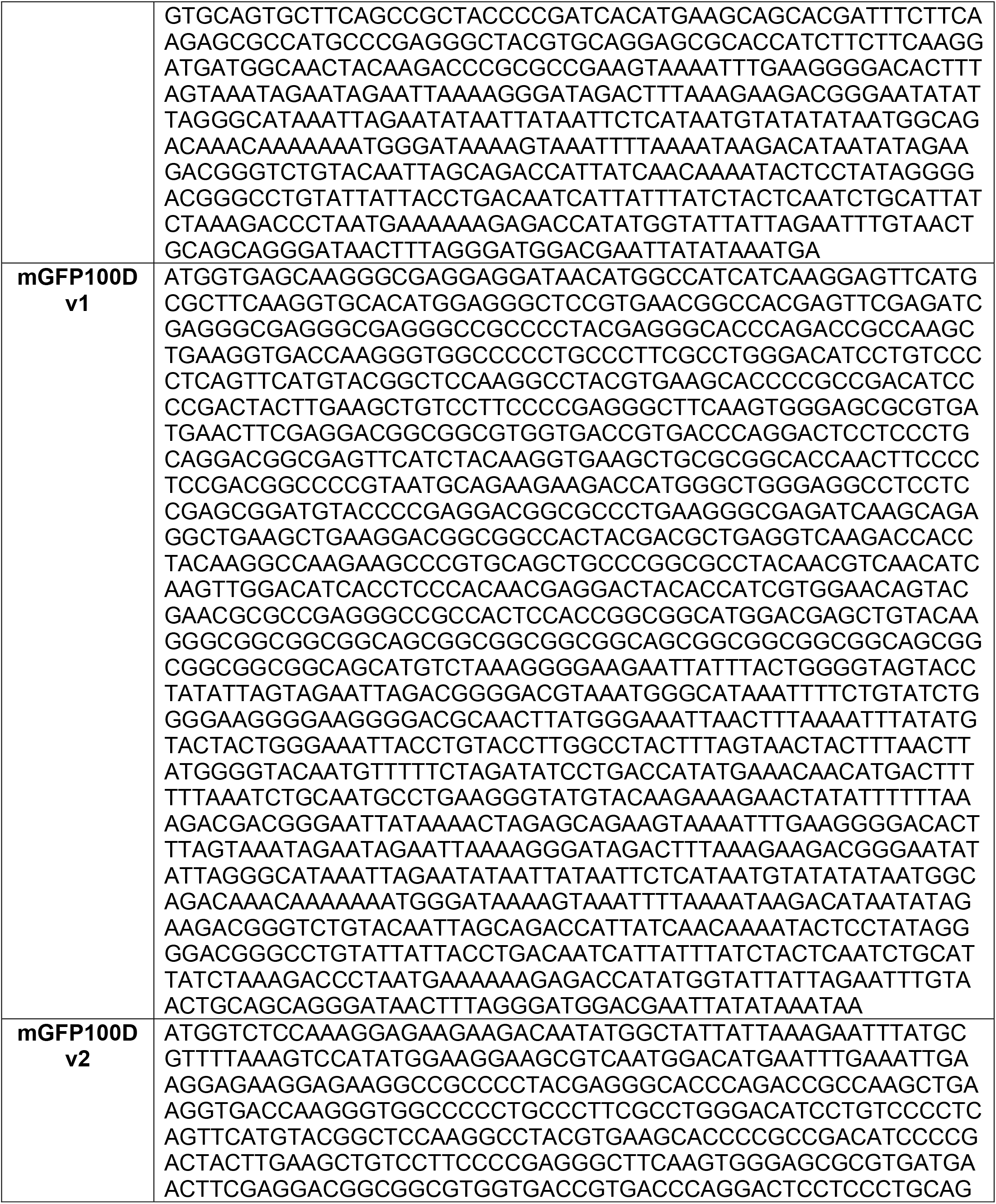
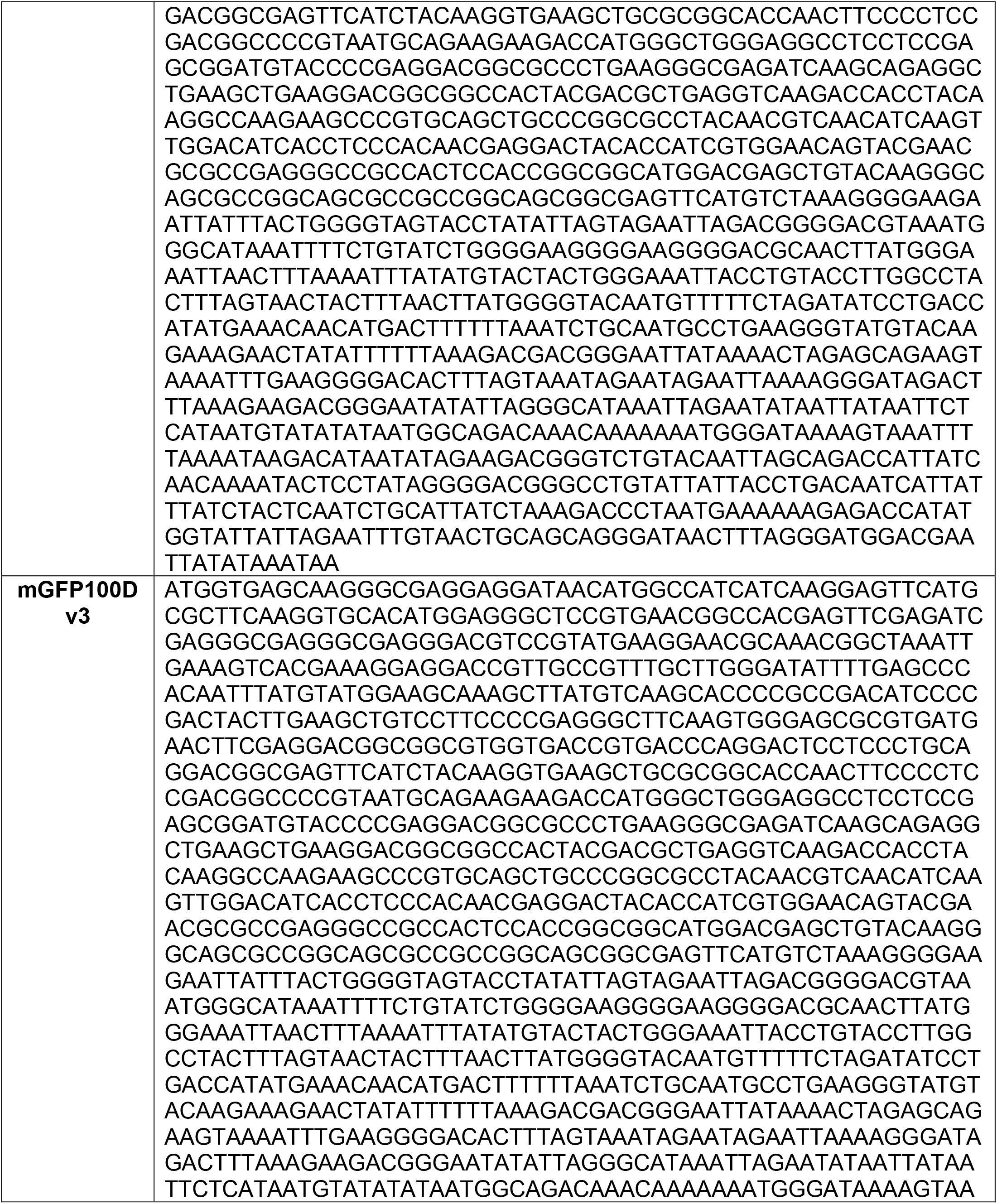
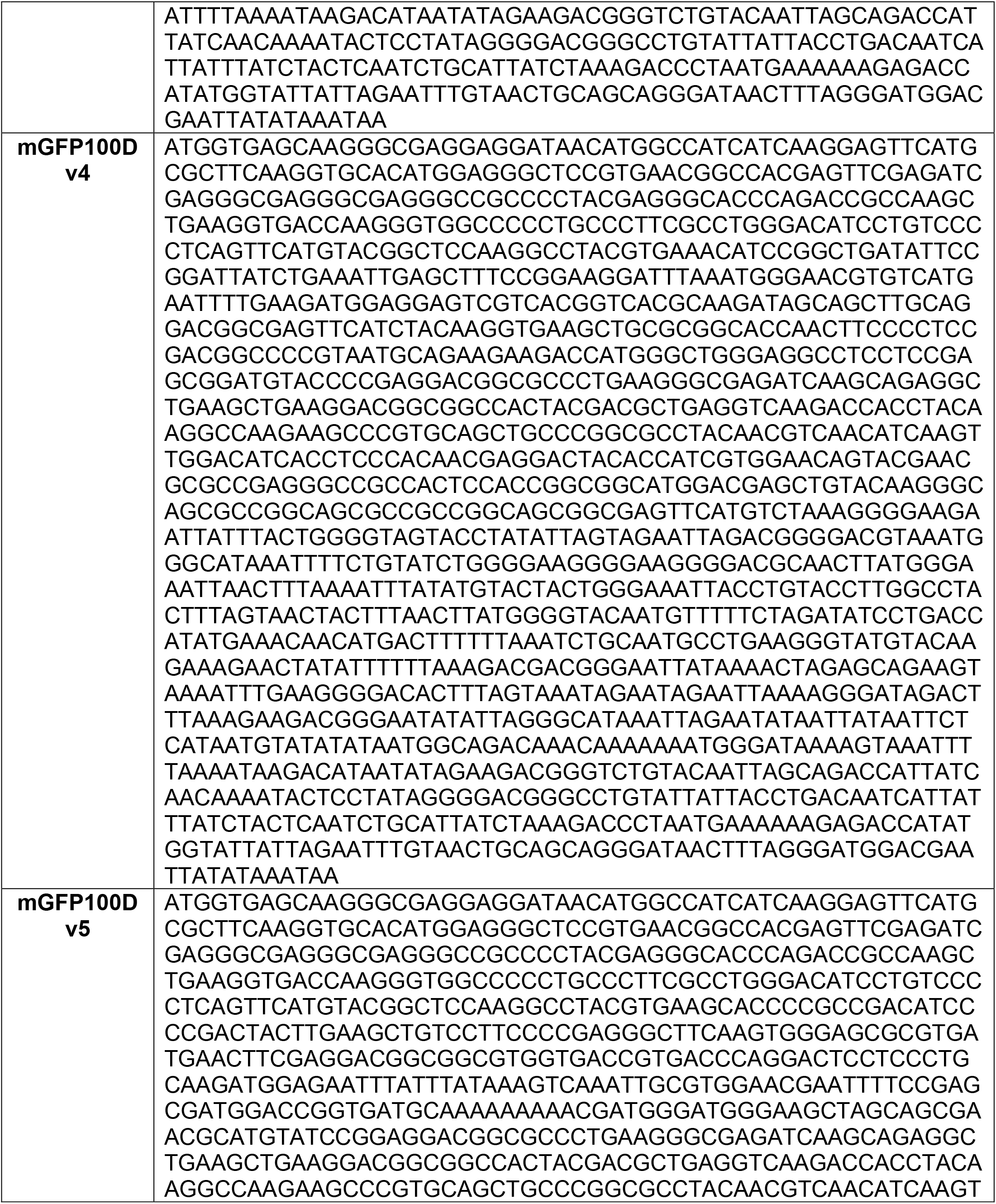
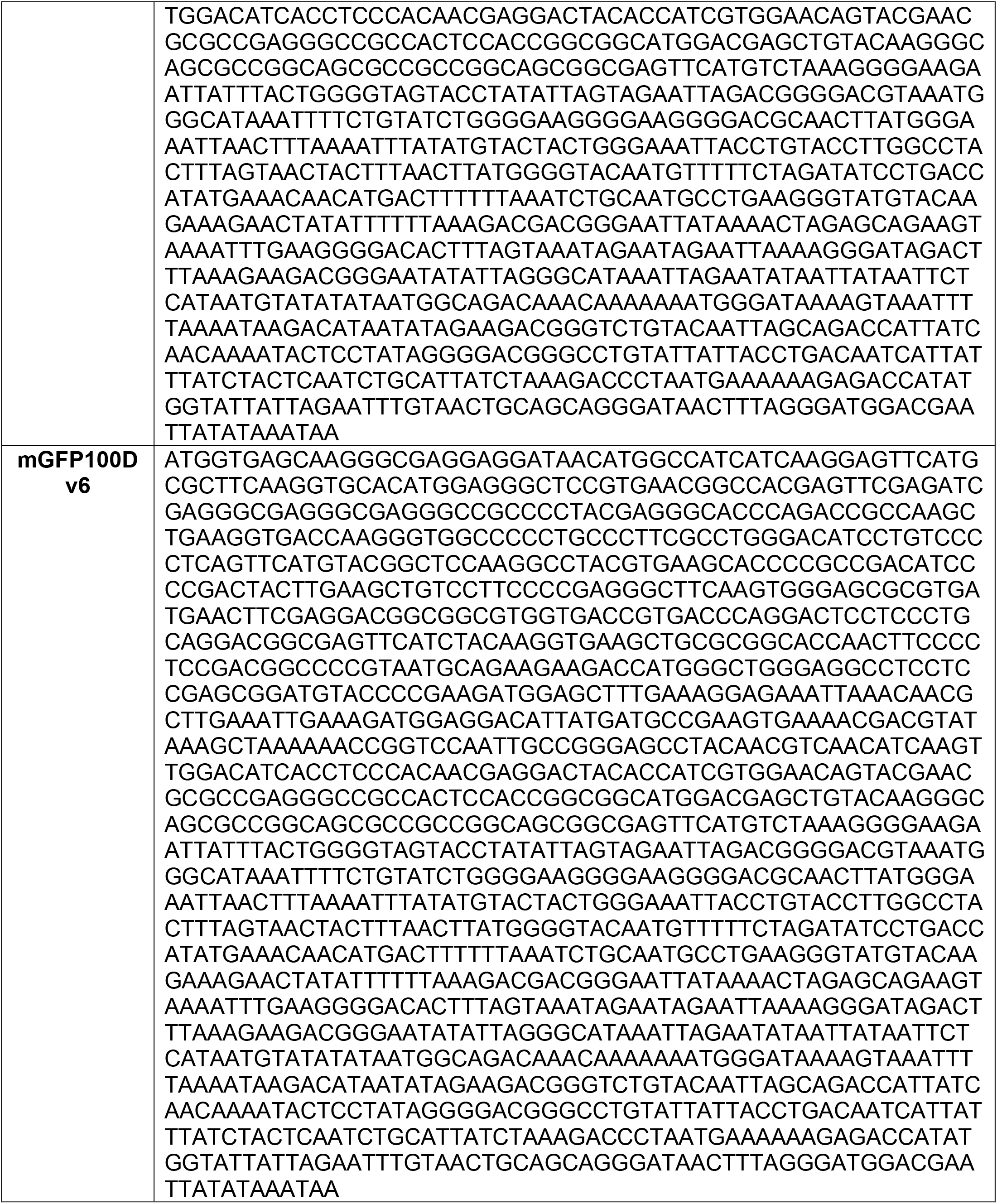
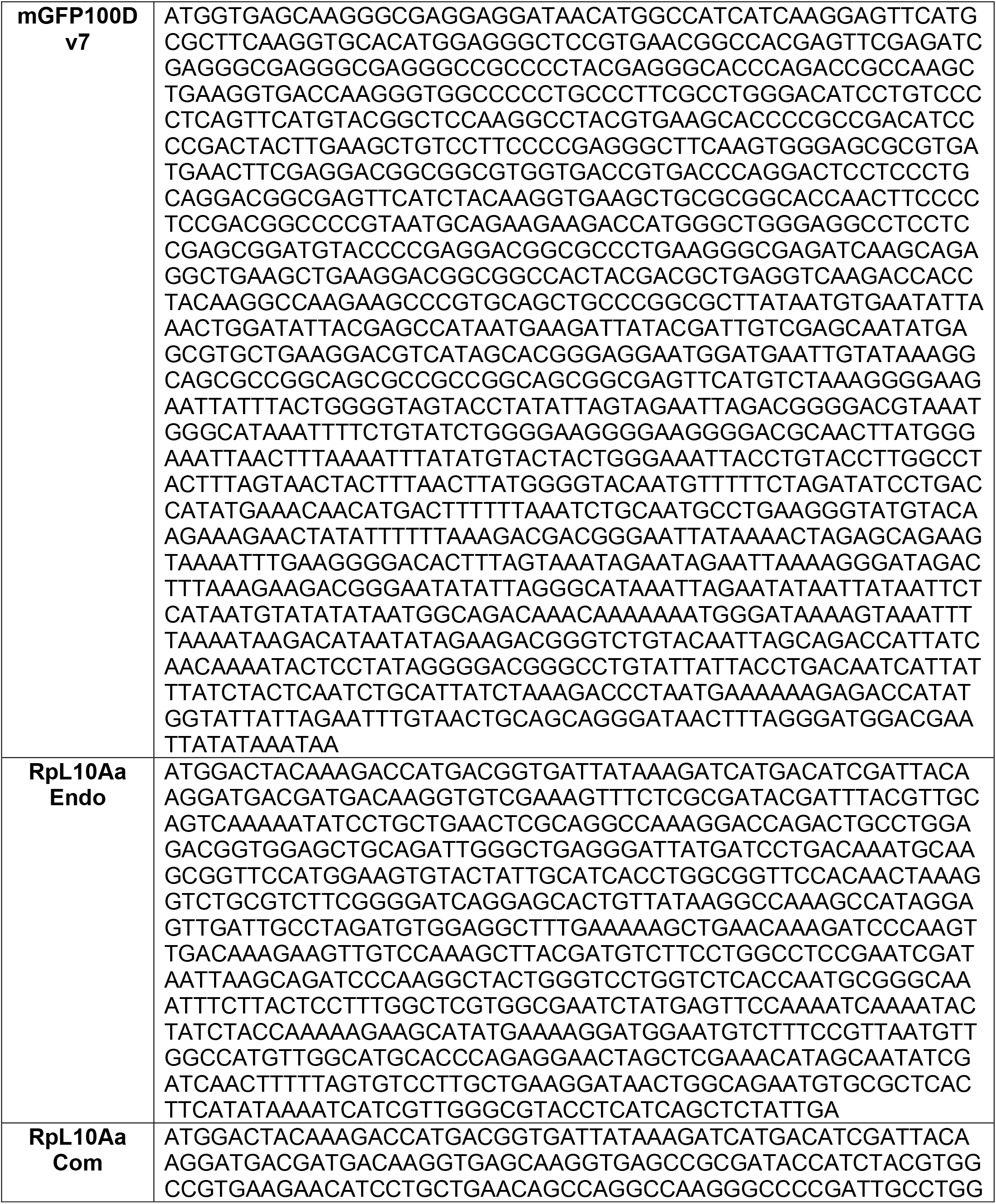
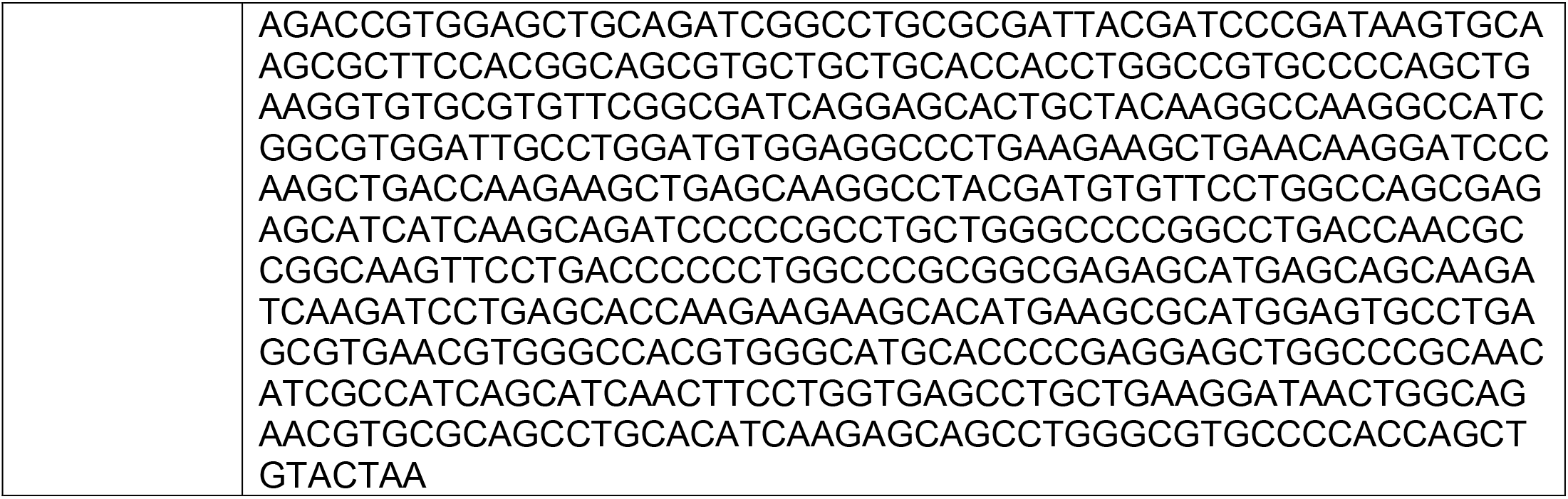
Full coding sequences of reporters.

**Table S2.** Occurrences of codons known to impair translation do not explain reporter expression patterns.

| **Reporter Name** | **Count CGA codons** | **Count AGG codons** | **Count AGG-AGG codon pairs** |
| --- | --- | --- | --- |
| **GFP0D** | 0 | 0 | 0 |
| **GFP30D** | 1 | 1 | 0 |
| **GFP50D** | 1 | 1 | 0 |
| **GFP60Dv1** | 0 | 1 | 0 |
| **GFP60Dv2** | 1 | 0 | 0 |
| **GFP60Dv3** | 1 | 0 | 0 |
| **GFP70D** | 1 | 0 | 0 |
| **GFP80D** | 0 | 0 | 0 |
| **GFP90D** | 0 | 0 | 0 |
| **GFP100D** | 0 | 0 | 0 |
| **GFP50C3'** | 0 | 0 | 0 |
| **GFP60C3'** | 0 | 0 | 0 |
| **GFP70C3'** | 0 | 0 | 0 |
| **GFP80C3'** | 0 | 0 | 0 |
| **GFP90C3'** | 0 | 0 | 0 |
| **GFP50C5'** | 0 | 1 | 0 |
| **GFP54C3'** | 0 | 0 | 0 |
| **mGFP100Dv1** | 0 | 1 | 0 |
| **mGFP100Dv2** | 0 | 1 | 0 |
| **mGFP100Dv3** | 0 | 1 | 0 |
| **mGFP100Dv4** | 0 | 1 | 0 |
| **mGFP100Dv5** | 0 | 1 | 0 |
| **mGFP100Dv6** | 0 | 0 | 0 |
| **mGFP100Dv7** | 0 | 1 | 0 |
| **RpL10Aa Endo** | 1 | 3 | 0 |
| **RpL10Aa Com** | 0 | 0 | 0 |

**Table S3.** Identity of child genes that gained max expression in the testis.

| **Child Gene** | **Highest Expressed in Testis?** | **Parent Gene** | **Highest Expressed in Testis?** |
| --- | --- | --- | --- |
| FBgn0030215 | TRUE | FBgn0036726 | FALSE |
| FBgn0038281 | TRUE | FBgn0036213 | FALSE |
| FBgn0035585 | TRUE | FBgn0016119 | FALSE |
| FBgn0051952 | TRUE | FBgn0265523 | FALSE |
| FBgn0034822 | TRUE | FBgn0030616 | FALSE |
| FBgn0031345 | TRUE | FBgn0030329 | FALSE |
| FBgn0261395 | TRUE | FBgn0261394 | FALSE |
| FBgn0034205 | TRUE | FBgn0031456 | FALSE |
| FBgn0036212 | TRUE | FBgn0023177 | FALSE |
| FBgn0029728 | TRUE | FBgn0030459 | FALSE |
| FBgn0028513 | TRUE | FBgn0010497 | FALSE |
| FBgn0052971 | TRUE | FBgn0040339 | FALSE |
| FBgn0051924 | TRUE | FBgn0019982 | FALSE |
| FBgn0036497 | TRUE | FBgn0020255 | FALSE |
| FBgn0265606 | TRUE | FBgn0004066 | FALSE |
| FBgn0015008 | TRUE | FBgn0025633 | FALSE |
| FBgn0036747 | TRUE | FBgn0031170 | FALSE |
| FBgn0034651 | TRUE | FBgn0264694 | FALSE |
| FBgn0039576 | TRUE | FBgn0030733 | FALSE |
| FBgn0050075 | TRUE | FBgn0003401 | FALSE |
| FBgn0032439 | TRUE | FBgn0010774 | FALSE |
| FBgn0053017 | TRUE | FBgn0035253 | FALSE |
| FBgn0034837 | TRUE | FBgn0015288 | FALSE |
| FBgn0250904 | TRUE | FBgn0033603 | FALSE |
| FBgn0030377 | TRUE | FBgn0015622 | FALSE |
| FBgn0039622 | TRUE | FBgn0015218 | FALSE |
| FBgn0038486 | TRUE | FBgn0037440 | FALSE |
| FBgn0030979 | TRUE | FBgn0036717 | FALSE |
| FBgn0053300 | TRUE | FBgn0052602 | FALSE |
| FBgn0036496 | TRUE | FBgn0025790 | FALSE |
| FBgn0052847 | TRUE | FBgn0030693 | FALSE |
| FBgn0051477 | TRUE | FBgn0014391 | FALSE |
| FBgn0264077 | TRUE | FBgn0015622 | FALSE |
| FBgn0034937 | TRUE | FBgn0262699 | FALSE |
| FBgn0039371 | TRUE | FBgn0033906 | FALSE |
| FBgn0038209 | TRUE | FBgn0013305 | FALSE |
| FBgn0015025 | TRUE | FBgn0039860 | FALSE |
| FBgn0015520 | TRUE | FBgn0004227 | FALSE |
| FBgn0032680 | TRUE | FBgn0031145 | FALSE |
| FBgn0031504 | TRUE | FBgn0039970 | FALSE |
| FBgn0031856 | TRUE | FBgn0024991 | FALSE |
| FBgn0052110 | TRUE | FBgn0027603 | FALSE |
| FBgn0030937 | TRUE | FBgn0034098 | FALSE |
| FBgn0050354 | TRUE | FBgn0260008 | FALSE |

## Notes

### Competing Interest Statement

The authors have declared no competing interest.

